# Long-term *in vivo* imaging of mouse spinal cord through an optically cleared intervertebral window

**DOI:** 10.1101/2021.09.14.460247

**Authors:** Wanjie Wu, Sicong He, Junqiang Wu, Congping Chen, Xuesong Li, Kai Liu, Jianan Y. Qu

## Abstract

Spinal cord, as part of the central nervous system, accounts for the main communication pathway between the brain and the peripheral nervous system. Spinal cord injury is a devastating and largely irreversible neurological trauma, and can result in lifelong disability and paralysis with no available cure. *In vivo* spinal cord imaging in mouse models without introducing immunological artifacts is critical to understand spinal cord pathology and discover effective treatments. We developed a minimal-invasive intervertebral window by retaining ligamentum flavum to protect the underlying spinal cord. By introducing an optical clearing method, we achieved repeated two-photon fluorescence and stimulated Raman scattering imaging at subcellular resolution with up to 16 imaging sessions over 167 days and observed no inflammatory response. Using this optically cleared intervertebral window, we studied the neuron-glia dynamics following laser axotomy and observed strengthened contact of microglia with the nodes of Ranvier during axonal degeneration. By enabling long-term, repetitive, stable, high-resolution and inflammation-free imaging of mouse spinal cord, our method provides a reliable platform in the research aiming at understanding and treatment of spinal cord pathology.

## 1. INTRODUCTION

*In vivo* imaging of the central nervous system (CNS) of small animal models is a crucial means of understanding the function of the CNS and its response to injury or diseases. In recent decades, nonlinear optical (NLO) microscopy has emerged as a powerful tool for the high-resolution imaging of biological tissues, including the CNS. Imaging of the live brain with sub-cellular resolution has been achieved using NLO microscopy through a cranial window in the mouse skull^1^. However, this open-skull procedure induces inflammation, indicated by microglia and astrocyte activation^2^, which can alter neuronal physiology^2^ and pia blood vessels^3^. To avoid the inflammation caused by surgery, thinned-skull protocols^4,5^ were developed, which provides a minimally invasive way to study cell dynamics in both healthy^6,7^ and pathological conditions^8,9^ in the living brain. As with the brain, imaging the spinal cord without inflammation induced by surgical preparation has been in high demand for spinal cord studies, including spinal cord injury, multiple sclerosis, neuropathic pain and spinal cord ischemia. However, preparing a spinal window in a mouse is much more challenging than a cranial window because of the more complex gross anatomy and large motion artifacts caused by the heartbeat and breathing.

To acquire high-quality optical images of the spinal cord, acute surgical preparation is usually adopted with a limited time window of several hours^10,11^. During preparation, the spinal cord is exposed by performing a dorsal laminectomy. Sometimes, dura is removed to increase imaging depth and artificial ventilation is used to minimize motion artifacts caused by breathing^12,13^. However, this procedure inevitably disturbs the spinal cord tissue and usually causes mild trauma. Furthermore, longitudinal imaging requires repetitive surgery and permits only up to six imaging sessions because of the increasing difficulty of repetitive surgery^10,13–15^. Another method of implanting a spinal chamber can achieve long-term imaging without the requirement of repetitive surgery^16–18^. However, a transient increase in the density of microglia and other inflammatory cells was observed after window implantation, because an immune response was activated in the spinal cord^16,17^. To increase window clarity and tolerance to implants, pharmacologic management of inflammation is required, which may affect the disease process being investigated. Recently, another protocol, spinal cord imaging through the intervertebral spaces without performing a dorsal laminectomy, has been proposed as a less invasive way to provide optical access to the spinal cord^19,20^. By removing muscle and ligament tissues between adjacent vertebrae, the spinal cord was imaged with only dura left. Using this protocol, it is reported that microglia activation was not observed by 2-hour time-lapse imaging after surgery, though clear microglia imaging and quantitative analysis were not demonstrated in the study^19^. Repetitive surgery coupled with an intervertebral window enabled longitudinal imaging with up to ten separate imaging sessions over more than 200 days^20^, which is comparable to the performance of a chronic implanted window^16^.

Despite the less invasive protocol of the intervertebral window, the inflammatory response to this surgical preparation has not been studied well, and it remains unclear whether an intervertebral window can serve as a reliable method to study neuroinflammatory disorders in the spinal cord without surgery-induced artifacts. In this work, we propose an improved intervertebral window protocol which retains the ligamentum flavum to significantly decrease the risk of activating microglia. In addition, to overcome the scattering issue induced by the ligamentum flavum and improve the image quality of the spinal cord, we adopted an optical clearing technique using a nontoxic chemical, Iodixanol, to treat the window interface. Using this method, we achieved subcellular-resolution, longitudinal imaging of the spinal cord with 16 imaging sessions over 167 days without an inflammatory response. With this minimally invasive long-term intervertebral window, we used a multimodal NLO microscope system (**Supplementary Fig. 1**) to study the neuron-glia dynamics following imaging-guided laser injury of axons. We further investigated the interaction between microglia and the nodes of Ranvier under normal and injured conditions. Different types of dynamic glia-node interaction were classified based on time-lapse imaging, and significantly strengthened contact between microglia and the nodes of Ranvier was observed after the distal axon was injured by laser ablation.

## 2. RESULTS

### 2.1 Intervertebral window for *in vivo* imaging of spinal cord

We investigated the behavior of microglia in the spinal cord of Cx3CR1-GFP mice following preparations of a conventional intervertebral window and a new intervertebral window of retaining ligamentum flavum, respectively. The microglial morphology was used as an indicator of inflammatory activity. It is known that microglial cells are the primary immune effector cells in the CNS. In the homeostatic state, microglia are highly ramified and dynamic, with their motile processes continually probing the tissue’s microenvironment^21–23^. On exposure to pathogen- or damage-associated molecular patterns, microglia are activated and change their morphology from ramified to amoeboid with enlarged soma and retracted processes^24–27^. As microglial phenotypes are inextricably associated with their function^28–31^, microglial morphology has been used widely as an objective criterion by which microglia activation and inflammatory activity in the CNS can be identified^30,32–40^. Notably, a number of studies have used a set of morphological parameters to describe the shapes of microglia cells and analyzed their dependence on the level of activation, which was assessed using immunohistochemical staining of cytokine signatures to highlight inflammatory activation^30,33,41–43^. Quantitative analysis showed that the morphology of microglia changes progressively with the level of expression of various inflammatory cytokines including IL-1β, IBA-1, CD11b, and CD68^30,33,41,43^. Unlike the immunostaining method that is only applicable to postmortem study, the morphological analysis of microglia combined with high-resolution *in vivo* imaging techniques can serve as a versatile and sensitive means to detect subtle inflammatory activity in living animals, which is indispensable for *in vivo* longitudinal study of immune responses to different pathological situations. Among the morphological parameters, the ramification index (RI)^32,44–47^ and the number of process endpoints (NPE)^33,36,48–50^ are widely used to describe the ramification of microglial cells quantitatively. RI is calculated as the ratio of the cell’s perimeter to its area normalized to that of a circle with the same area^32^, while NPE counts the total number of microglial cell processes^33^. Significant decreases in both the ramification index and the endpoints of microglia are typical symptoms of high degrees of inflammatory activation, which is evident in different pathological models of neuroinflammation such as diffuse brain injury, ischemic stroke, peripheral nerve injury, etc^33,44,47,50^.

To study the microglial phenotypes under different conditions, mice were divided into three groups (three mice per group) (**Supplementary Fig. 2**). The first group of mice had dorsal column crush (DCC) injury performed at the T12 level after a laminectomy to generate an acute inflammatory process, so that the morphology of activated microglia can be characterized first as a positive control. The second group of mice underwent the conventional surgical procedure of intervertebral window preparation to expose the spinal cord in the intervertebral gap. One hour after surgery, mice in these two groups were imaged by a two-photon excited fluorescence (TPEF) microscope for two hours and the behavior of microglia was recorded over a 30-min interval. After imaging, all the mice were perfused for histological analysis. In addition, the third group of mice, which did not undergo any surgery before the histological study, were used as a negative control. Next, we examined the microglia morphology in fixed spinal cord slices of the three groups of mice (**Supplementary Fig. 3a**). The microglial cells in the region of 0–50 *μ*m below the dorsal surface were selected for analysis, corresponding to the *in vivo* imaging depth. By comparison with the negative control group, the small values of RI and NPE in the positive control group indicates severe activation of microglia after spinal cord injury (**Supplementary Fig. 3b**,**c**). Of the three mice with intervertebral windows, one mouse (#2) showed significantly decreased RI and NPE and aggregation of microglia was also found close to the dorsal surface (**Supplementary Fig. 3a**), while the other two mice showed comparable morphological indices to the negative control group. Then we compared the *in vivo* results (**Supplementary Fig. 3d-f**) with those of histopathology studies. The *in vivo* time-lapse imaging shows that microglia in the spinal cord injured by DCC were activated with significantly decreased ramification at the beginning of imaging and little change over the following two hours (**Supplementary Fig. 3e**,**f**). This suggests that microglia can respond quickly to pathological insults and be activated within an hour. In the mice with an intervertebral window, microglia showed differentiated but stable ramification during the two-hour observation. Consistently with histological results, the mouse (#2) with activated microglia in histological analysis also showed activation of microglia with retraction of fine processes in *in vivo* studies (**Supplementary Fig. 3e**,**f**). To evaluate rigorously the possibility of the activation of microglia during longitudinal imaging following surgical preparation of an intervertebral window, we repeatedly imaged 12 mice with 2-6 day intervals between the adjacent imaging sessions. The results show that activation of microglia was found in 58% (7/12) of the mice in the first imaging session, and 75% (3/4) of the mice in the second imaging session (**Supplementary Fig. 3h**). None of the mice underwent surgery three times without activation of microglia (**Supplementary Fig. 3h-k**). Notably, the difficulty of surgery increased significantly in later procedures because of the growth of scar tissue adhering to the surface of the spinal cord. This result suggests that intervertebral window preparation will inevitably cause irritation to the spinal cord and induce activation of microglia, preventing the inflammation-free longitudinal study of the spinal cord.

Prompted by the thinned-skull procedure, we explored whether we can lower the risk of inflammation by retaining the ligamentum flavum during surgical preparation of the intervertebral window (**Supplementary Fig. 4**). The ligamentum flavum (LF) is a series of ligaments composed of elastic fibers and collagen. They join the laminae of the adjacent vertebra and are located directly above the spinal cord from a posterosuperior view, separated by the meninges and epidural space (**Fig. 1a,b**)^51,52^. The epidural space contains adipose tissue and blood vessels, which, together with ligamentum flavum, protects the underlying spinal cord, but makes the whole window optically inhomogeneous and less transparent (**Fig. 1c-e**). A small number of cells labeled by Texas Red Dextran above the spinal cord are probably phagocytic immune cells, which is also observed in previous studies^16,17,53^ (**Fig. 1e**). Nevertheless, we found that high-resolution images of the spinal cord can still be captured in a small field of view (FOV) without there being adipose tissue and blood vessels along the optical path (**Supplementary Fig. 5**).

**Figure 1.**
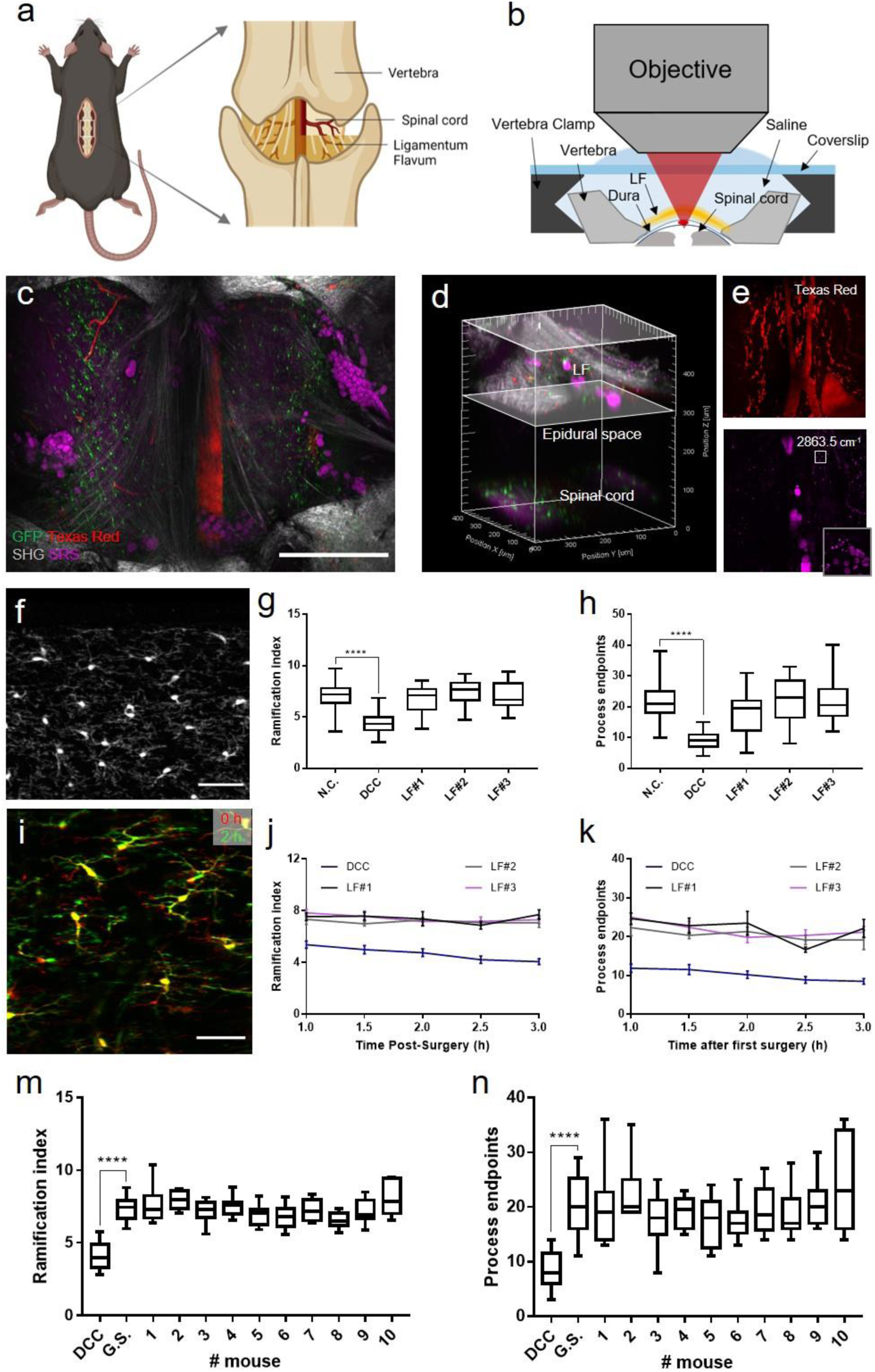
Intervertebral window of retaining ligamentum flavum. (a) Schematic diagram of the intervertebral window with ligamentum flavum (LF). (b) Cross-sectional schematics of the LF window. (c, d) Projection and 3D reconstruction of a multimodal image stack through the LF window to spinal cord surface in a live Cx3CR1 transgenic mouse. Green: GFP labeled microglia; Red: blood vessels labeled with Texas Red dextran; Gray: second harmonic generation (SHG) signals of collagen and other connective tissues; Magenta: stimulated Raman scattering (SRS) signals of adipose tissue and myelin with the Raman shift at 2863.5 cm^-1^, attributed to the vibration of the methylene group enriched in lipids. Scale bar, 500 *μ*m. (e) Maximal projection of the image volume between the two planes of depth from 300-500 *μ*m indicated in (d) showing the distribution of blood vessels (Texas Red dextran in the red fluorescence channel) and adipocytes (SRS imaging channel) in the epidural space. Cells labeled by Texas Red are probably invading immune cells. Cells shown in the inset with strong pump-probe absorption at 2863.5 cm^-1^ are red blood cells indicated by their specific dumbbell shape. Scale bar, 50 *μ*m. (f) Two-photon fluorescence image of a 50-*μ*m-thick longitudinal spinal cord slice under the LF window. Scale bar, 50 *μ*m. (g, h) Evaluation of the microglial ramification index (g) and number of process endpoints (h) of spinal cord fixed slices from the LF window group, the dorsal column crush (DCC) group and the negative control (N.C.) group. The boxplots are shown with median, upper and lower quartiles and maximum and minimum values. Kruskal-Wallis test: ****P < 0.0001, n ≥ 20 measurements from 6-8 slices per mouse, three mice per group. (i) The *in vivo* superimposed images of microglia at an interval of two hours, showing ramified microglia morphology with highly motile processes under the LF window. Scale bar: 50 *μ*m. (j, k) Changes of the microglia ramification index (j) and number of process endpoints (k) during two-hour *in vivo* imaging in the LF window group and the DCC group. n ≥ 6 measurements per time point per mouse. Error bars, s.e.m. (m, n) *In vivo* evaluation of the microglial ramification index (m) and process endpoints (n) of ten mice with LF window at the first live imaging session. The *in vivo* morphological indices from the three non-activated mice with LF (LF#1-3 in (g, h)) were used as the gold standard (G.S.) for *in vivo* microglia activation evaluation. The *in vivo* results from the DCC group were used as the positive control. Microglia activation in each mice was determined by comparing the calculated ramification index and number of process endpoints with the G.S. Kruskal-Wallis test: ****P < 0.0001, n ≥ 6 microglial cells for morphological quantification for each mouse; the box plots are shown with median, upper and lower quartiles and max and min values.

To evaluate the activation of microglia beneath the new intervertebral LF window, we characterized the morphology of microglia both *in vivo* and in fixed spinal cord using high-resolution TPEF imaging, and compared it to that of intact and injured spinal cords. Histological results show that microglia under the window showed ramified morphology with similar RI and NPE to the negative control group **(Fig.1f-h)**. Meanwhile, *in vivo* time-lapse imaging suggests that microglia retained ramified morphology during the two-hour imaging period **(Fig. 1i-k)**. To validate the repeatability of the surgical preparation of the new window, we imaged another ten mice through the LF window and conducted quantitative morphological analysis of microglia. Notably, we found none of the mice showed activation of microglia (**Fig. 1m,n**). This result indicates that retaining LF can indeed prevent microglia activation, and thus this protocol of LF window can serve as a minimally invasive method for *in vivo* optical imaging of the spinal cord.

### 2.2 Optical clearing intervertebral window of retaining ligamentum flavum

Although retaining LF helps to reduce the risk of inflammation caused by surgery, the LF layer as well as tissues in the epidural space introduces optical scattering to decrease the penetration depth of spinal cord imaging. After initial surgery, both LF and the epidural space were infiltrated and filled by a large number of cells, which greatly decreased image contrast and resolution (**Supplementary Fig. 6**). In this work, optical clearing method was developed to reduce the optical inhomogeneity of the LF window. Recently, Iodixanol, an FDA approved non-toxic compound commonly used as a contrast agent during coronary angiography, has been shown to improve image quality by refractive index matching in live specimens^54^. We tested the applicability of Iodixanol as an optical clearing medium to facilitate *in vivo* spinal cord imaging through the LF window. On day1 post-surgery when cell infiltration reduced the transparency of the window, we applied Iodixanol on top of the LF layer and found a significant improvement in the window transparency and optical homogeneity after 10 minutes (**Fig. 2a,b**). The application of Iodixanol restored the image contrast and resolution of both two-photon and stimulated Raman scattering (SRS) imaging on day 1 to almost the same level as on day 0 (**Fig. 2c-e**). Importantly, this improvement can be lost by replacing the Iodixanol with saline (**Fig. 2f**) and then recovered by reapplying Iodixanol (**Fig. 2g**). This phenomenon indicates that the reduction in optical inhomogeneity should be achieved by refractive index matching rather than by direct removal of scatterers in tissues. Multimodal imaging of the spinal cord with epidural space and ligamentum flavum showed the tissue structure to be consistent before and after Iodixanol application (**Fig. 2e-g**), which further confirms our hypothesis concerning the optical clearing mechanism of Iodixanol. By increasing the concentration of Iodixanol up to 60% w/v (*n* ≈*1*.*429*), the improvement in imaging increased further, indicating better matching of refractive indices (**Supplementary Fig. 7**).

**Fig 2.**
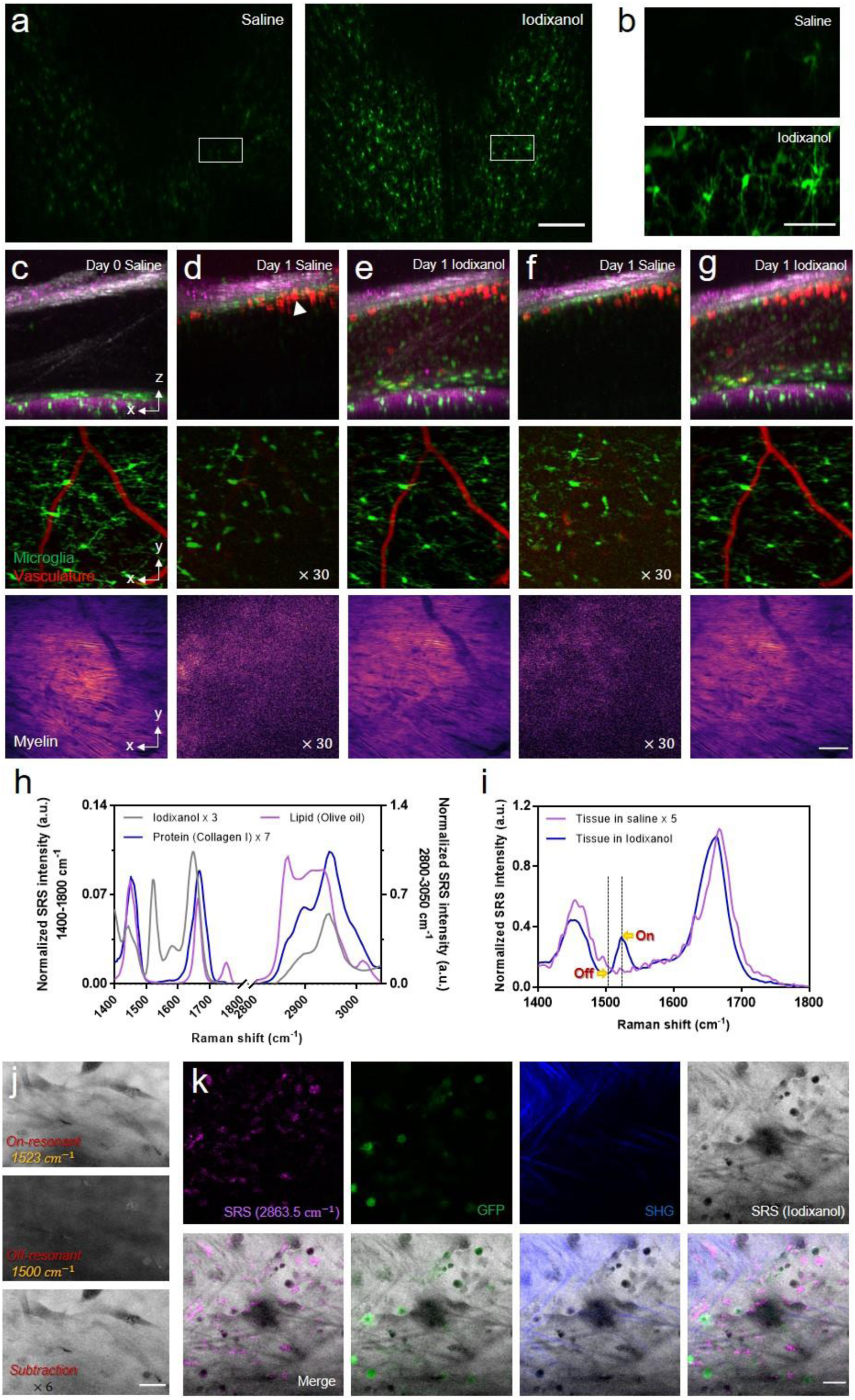
Optical clearance of intervertebral LF window by using Iodixanol. (a) Maximal projection of a microglia TPEF image stack under LF window before and after optical clearing on day 1. Scale bar, 200 *μ*m. (b) Magnification of the box region in (a) shows the detailed microglia image before and after optical clearing. Scale bar, 50 *μ*m. (c-g) Top row: Maximal x-z projection of the multimodal images of the LF window. Green, GFP-labeled microglia and other immune cells; Gray: SHG of connective tissues; Red: Texas Red labeled blood vessels and invading, likely inflammatory cells (white arrowhead); Magenta, SRS of lipid at Raman shift of 2863.5 cm^-1^; Middle row: the x-y maximal TPEF projection images of microglia (green) and vasculature (red) under LF window; Bottom row: the x-y maximal SRS projection images of myelin under LF window. The images were captured on day 0 (c) and day1 (d-g) after first surgery. On day 1, the window was firstly immersed in saline (d) and then replaced with Iodixanol (e) which was removed (f) and reapplied (g) again at later times to verify the repeatability of the clearing effect. All the x-z and x-y projection images were normalized to the same value for each imaging modality. The signal intensity in x-y projection images of (d) and (f) was digitally enhanced by 30 times for better visualization. Scale bar, 50 *μ*m. (h) The SRS spectra of Iodixanol (60% w/v), protein (Type I collagen) and lipid (Olive oil) at the fingerprint (1400-1800 cm^-1^) and Carbon-Hydrogen stretching region (2800-3150 cm^-1^). The spectral intensity was subtracted by the non-SRS background with the flat spectral response and was normalized by the lipid CH_2_ peak at 2863.5 cm^-1^. The SRS spectral intensity of Iodixanol and protein were digitally enhanced 3 and 7 times for better visualization. (i) On day 9, SRS spectra of the LF layer immersed in saline and 60% w/v Iodixanol, respectively. The two spectra were normalized by the peak intensity value of the Iodixanol immersed tissue at a vibrational frequency of 1663.3 cm^-1^. The SRS spectral intensity of tissue immersed in saline was digitally enhanced 5 times for better visualization. (j) The SRS image of the LF layer immersed in 60% Iodixanol at 1523 cm^-1^ (on-resonant) and 1500 cm^-1^ (off-resonant) Raman shift and their subtraction. Scale bar, 20 *μ*m. (k) The *in vivo* multimodal NLO images of the Iodixanol immersed LF layer on day 2 showing the distribution of Iodixanol in the interstitial space. Scale bar, 20 *μ*m.

Next, we investigated how refractive index matching was achieved in the LF window by using hyperspectral SRS imaging combined with two-photon microscopy. We first acquired the SRS spectrum of Iodixanol in the fingerprint region (1400-1800 cm^-1^) as well as in the Carbon-Hydrogen (C-H) stretching region (2800-3150 cm^-1^) and compared it with typical spectra of lipid (olive oil) and protein (type I collagen) (**Fig. 2h**). Iodixanol shows a similar spectrum to protein from 2800 to 3150 cm^-1^, but has a quite different spectrum profile in the fingerprint region where there is a unique vibrational peak at 1523 cm^-1^ contributed by the aromatic ring as well as the secondary amide II band in its molecular structure^55^. By sweeping the SRS spectrum of the LF layer immersed in Iodixanol, we found a small SRS peak at 1523 cm^-1^ that disappeared when Iodixanol was rinsed out (**Fig. 2i**). Therefore, 1523 cm^-1^ is a vibrational peak contributed solely by Iodxianol, which can be used to visualize the Iodixanol distribution without interference from other endogenous biomolecules. To eliminate non-Raman background interference, a subtraction method was used to obtain the genuine SRS signal of Iodxianol (*I*_SRS_ = *I*_ON_ - *I*_OFF_)^56,57^. Briefly, the SRS baseline signal (off-resonant, *I*_OFF_) at 1500 cm^-1^ was subtracted from the SRS peak signal (on-resonant, *I*_ON_) at 1523 cm^-1^ to suppress non-Raman backgrounds (**Fig. 2j**). To investigate the distribution of Iodixanol through the LF window, we applied multimodal NLO microscopy combing SRS, TPEF and second harmonic generation (SHG) to visualize Iodixanol, cells and collagens simultaneously. The multimodal images showed that collagen and cellular structures are spatially correlated with the negative contrast regions in the Iodixanol SRS images (*I*_SRS_) (**Fig. 2k**), indicating that Iodixanol achieved refractive index matching primarily by increasing the refractive index of the interstitial fluid. The clearing effect of Iodixanol becomes worse with time due to the gradual dilution of the Iodixanol indicated by the decreased SRS signal of Iodixanol between the window surface and coverslip (**Supplementary Fig. 8, Supplementary Video 1**). Therefore, Iodixanol was supplemented hourly to maintain good refractive index matching, and the imaging was usually started 10 min after every Iodixanol administration when its optical clearing effect reached a plateau (**Supplementary Fig. 8**).

Safety issues are a crucial concern when applying optical clearing agent to living animals. Because of non-toxicity, Iodixanol has long been used as an intravenous X-ray contrast agent^58,59^ as well as a density gradient medium for cell isolation^60^. When applied as a refractive index matching media for live imaging, Iodixanol doesn’t show any toxic effects on living Hela cells, planarians and zebrafish^54^. In this study, we assessed the effects of exposing the spinal cord to Iodixanol by exploring microglia activation after optical clearing. We imaged microglia through the LF window before and 1hr after applying Iodixanol (60% w/v) on day 0 when high-resolution microglia images still could be acquired without optical clearing. Results show that microglia remained ramified and continually surveying the microenvironment with highly motile processes after Iodixanol administration (**Supplementary Fig. 9**). To further assess the potential long-term effects of exposing the spinal cord to Iodixanol, we continued to image microglia on day 1 and day 3 with the window treated with Iodixanol (60% w/v). Time-lapse *in vivo* imaging shows that all the microglial cells in the FOV retained ramified morphology with dynamic processes, indicating no inflammation (**Supplementary Fig. 9**). Collectively, these results demonstrate that applying Iodixanol to the intervertebral window does not impact the spinal cord, largely alleviating safety concerns.

Since optical clearing by Iodixanol can significantly increase the window clarity without activating microglia, we next explored the potential of this optical cleared intervertebral window for minimally invasive longitudinal imaging. We conducted time-lapse multimodal NLO imaging of four Cx3CR1-GFP mice through the window and achieved up to 16 imaging sessions over 167 days without observing microglia activation (**Fig. 3**). It was found that on day 0, optical clearing significantly increased the transparency and optical homogeneity of the whole window (**Supplementary Fig. 10**). Within the first week after initial surgery, scar tissue at the surgical site has not developed fully and the large interstitial space below the LF layer allowed easy matching of refractive indices by Iodixanol and therefore permitted high-resolution fluorescence imaging **(Supplementary Fig. 11)**. The improvement of fluorescence and SRS signal by optical clearing on day 4 reached about 20 times **(Supplementary Fig. 11c-f)**. Usually a week later, scar tissue developing with collagen, blood vessels, and recruited dense cells, severely degraded the window transparency and reduced the optical clearing effect (**Supplementary Fig. 11,12**). Therefore, it is necessary to remove the newly grown tissue above the LF layer. Due to the mechanical toughness of the LF layer, the loose granulation tissue at the top of the window can be easily distinguished. The precise surgical removal of scar tissue leaving the LF layer intact can be achieved with a high success rate. After tissue removal and Iodixanol treatment, spinal cord images with subcellular resolution could be recovered (**Supplementary Fig. 12**). During each imaging session, to equilibrate the heterogeneous refractive indices, Iodixanol was applied to the surface of the intervertebral window prior to NLO imaging. Although the structure of the LF window varies with time, optimal refractive index matching was always reached at about 10 min after Iodixanol treatment (**Supplementary Fig. 11f**). Therefore, Iodixanol was supplemented every hour and imaging was usually started 10 min after every Iodixanol administration.

**Figure 3.**
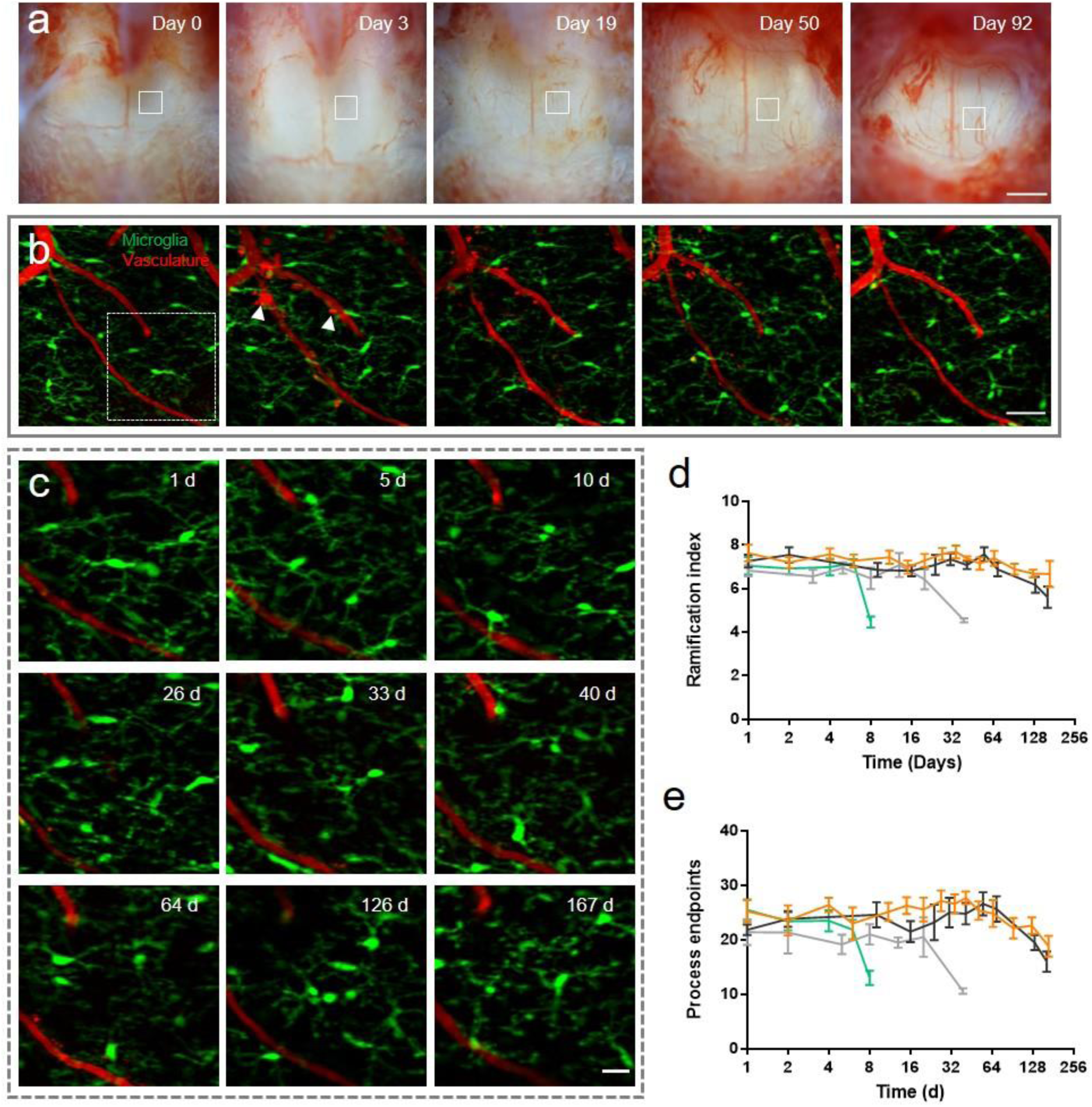
Long-term spinal cord imaging through optically cleared LF windows. (a) Bright-field images of an LF window over three months. Scale bar, 500 *μ*m. (b) Maximal projections of microglia (green) and vasculature (red) image stacks at the same site in the box region of (a). Vasculature labeled by Texas Red dextran was used to navigate to the same region of interest in different imaging sessions. The arrowheads indicate Texas Red dextran labeled perivascular cells and invading, likely inflammatory cells above the spinal cord. Scale bar, 50 *μ*m. (c) Magnification of the box region in (b) shows the detailed structures of microglia (green) and vasculature (red) at indicated times. Scale bar, 50 *μ* m. (d, e) Ramification index (d) and number of process endpoints (e) as functions of time during longitudinal two-photon fluorescence imaging. Each curve represents the statistical data of one mouse. For statistics of ramification index and number of process endpoints, 6-10 microglial cells with intact and clear morphology (contrast >0.97) were analyzed at each time point. The longitudinal study was terminated when microglial activation was observed, or the imaging region of interest was lost because of the shrunken field of view. Error bars, s.e.m.

We evaluated the inflammation response by morphological analysis of microglia in each imaging session. Microglial cells in the same region of interest (ROI) with intact and clear morphology (contrast > 0.97) (**Supplementary Fig.13**) were selected for morphological quantification The statistics of RI and NPE show that microglia activation with significantly decreased ramification was observed on day 8, day 39 and day 161 in three mice (**Fig. 3d,e**), respectively, due to accidental touch to the spinal cord by surgery tools when scar tissue removal was required. One mouse of four was imaged for as long as 167 days without inflammation (**Fig. 3a-c**), but the ROI was lost on day 203 because of the decreased FOV of the intervertebral window. From the bright-field images of the two mice which were imaged for more than 160 days, we observed the intervertebral window becoming smaller over time because of the growth of the surrounding rigid connective tissues (**Supplementary Fig. 14**), which limits the time span of the intervertebral window.

### 2.3 Multimodal NLO imaging of axonal degeneration after laser axotomy

Using double transgenic mice expressing EYFP in dorsal root ganglion afferent neurons and EGFP in microglia, we evaluated the response of axons and microglia to laser-induced injury. Axons and surrounding myelin sheaths were imaged together with microglia using our multimodal NLO microscope. YFP and GFP signals were differentiated using a spectral unmixing method as previously described^61^. We conducted precise single axon axotomy using tightly focused femtosecond laser pulses (**Fig.4, Supplementary Fig. 15**). Microglia responded rapidly to the injury by extending their cytoplasmic processes towards the lesion (**Supplementary Fig. 15b, Supplementary video 2**). The proximal end of the axon underwent acute degeneration within an hour post injury, and the surrounding myelin sheath kept close contact with the axon during degeneration (**Supplementary Fig. 15c**). After 1 day, dieback of the proximal ends slowed down and a large number of microglia as well as bone marrow-derived macrophages (BMDMs)^62^ were recruited to the injury site. Thanks to the precise laser axotomy on a single axon and high-resolution *in vivo* fluorescence imaging, we could observe clearly the spatially confined microglia/macrophage distribution strictly along the axonal degeneration path (**Fig. 4a**). Though the influx of microglia and BMDMs has been shown to correlate with axonal dieback^62,63^, our imaging-guided laser microsurgery along with longitudinal imaging permits study of the interaction between microglia/macrophages and injured axons in a much higher resolution both temporally and spatially. The results show that 1 day post injury (dpi), microglia mainly aggregated at the lesion site. At 3 dpi, however, the microglial aggregation moved along the direction of axon degeneration and kept physical contact with the proximal end of the injured axon. At 8 dpi, the aggregation disappeared and microglia were redistributed homogeneously in the FOV (**Fig. 4a-b**). This spatiotemporal distribution of microglia/macrophages could be correlated with its cellular function of tissue debris clearance. At 3 dpi, microglia/macrophages phagocytosis of the myelin and axon debris along the axonal degeneration path was observed (**Fig. 4c**). Our time-lapse multimodal imaging showed that the amount of myelin debris was significantly reduced, corresponding with the decreased density of microglia/macrophage at 8 dpi, with only a few debris left inside the cell bodies of myelin-laden microglia/macrophages (**Fig. 4d**). These results provide crucial *in vivo* evidence to support previous studies that observed microglia/macrophage engulfment of axon and myelin debris based on postmortem analysis^64,65^.

**Figure 4.**
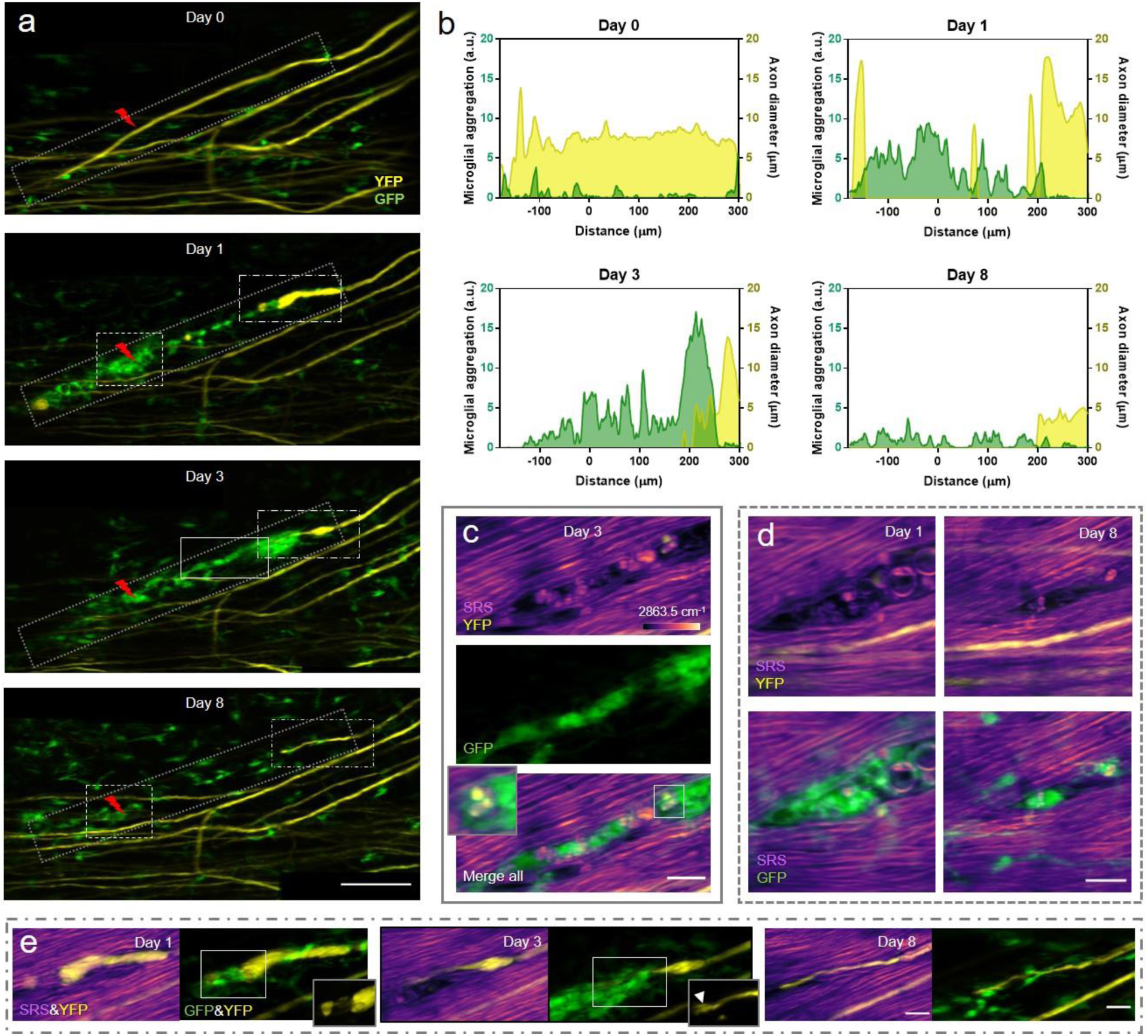
*In vivo* multimodal NLO imaging of axonal degeneration after laser axotomy. (a) Maximal z intensity projections of TPEF image stacks of YFP labeled axons (yellow) and GFP labeled microglia (green) at indicated times before and after laser axotomy. The lightning bolt symbol indicates the lesion site. Scale bar, 100 *μ*m. (b) The dynamics of the distribution of microglia (green) along the axonal degeneration path and the diameter of the injured axon (yellow). Only microglial cells located in the dot rectangular region along the degenerating axon in (a) were included for analysis. (c) The multimodal image of the spinal cord in the solid box region in (a) shows resident microglia and/or recruited macrophage aggregation along the axonal degeneration path at 3 days post injury (dpi). Colocalization of axon (yellow) and myelin (magenta) debris and microglia (green) indicates phagocytosis of microglia/macrophages. Insets, a zoomed-in view of myelin and axon debris colocalized with microglia/macrophage. (d) The multimodal images taken at 1dpi and 8 dpi in the dashed box region in (a) show the initialization and finalization of debris clearance, respectively. (e) The zoomed-in multimodal images of the axonal proximal end at indicated time points. The imaging area corresponds to the long dash-dot box region in (a). For clear visualization, the merged SRS and YFP images are shown as a single slice, while the merged GFP and YFP images are shown as the maximum z projections of image stacks. Insets, YFP images of the axonal proximal end. The arrowhead denotes small sprouts emerging from the tip of the axon. SRS images of myelin were taken at Raman shift of 2863.5 cm^-1^. Scale bars in (c-e), 20 *μ*m.

In addition to the phagocytosis of axon and myelin debris, microglia and macrophages were also reported to mediate axonal dieback by forming cell to cell contacts with the dystrophic endings of injured axons^62,63,66^. At 1 dpi, the injured axon formed an enlarged endbulb where the surrounding myelin sheath was lost (**Fig. 4e**). Interestingly, an axonal fragment was loosely connected to the enlarged proximal ends and surrounded by microglia/macrophages (**Fig. 4e**). It looks like the microglia/macrophages were pulling and stretching the fragment from the proximal end, as suspected in a previous study^63^. On the third day after injury, even stronger physical contact was observed between the microglia/macrophages and the proximal axonal end. Despite closely contacted by microglia/macrophages, the injured axon had limited secondary degeneration after day 1, and conversely, it showed early signs of regeneration. At 3 and 8 dpi, the dystrophic proximal ends became thinner and exhibited growth cone like structures with a small regeneration length of 12 *μ*m from 3 to 8 dpi. (**Fig. 4e**).

In addition, we also investigated microglial behavior and microglia-axon interaction following severe spinal cord injury by inflicting injury by laser over a large area (**Supplementary Fig. 16**). Large macrophage aggregations were observed at the lesion site at 1 dpi and expanded further in the following days. Compared with the phenomena observed in the single axon injury, the strict spatial correlation between the microglia/macrophage distribution and individual axonal degeneration path was not observed, probably because of the large size of the injury. In addition, the macrophage aggregation remained at the lesion site for at least a month, suggesting long-lasting inflammatory activity. The injured axon underwent acute and subacute degeneration within the first three days and then became almost immobilized in the following weeks. As we observed previously, the axonal ends first became enlarged and then thinned with sprouts appearance at 3 and 8 dpi. As can be seen, by taking advantage of the multimodal NLO imaging with high spatiotemporal resolution, we demonstrated a reliable model to study the highly dynamic processes of debris clearance and glia-neuron interactions during tissue injury and remodeling under finely controlled injury conditions.

### 2.4 Dynamic interaction between microglia and the nodes of Ranvier

Nodes of Ranvier, known as myelin-sheath gaps, are characterized by short and periodic regions of the axonal membrane that are bare of myelin^67,68^. The axolemma at nodes of Ranvier is exposed directly to the extracellular matrix and is highly enriched in ion channels, which permit the rapid exchange of ions to regenerate the action potential^68^. Therefore, nodes of Ranvier play a key role in fast saltatory propagation of action potentials. In the CNS, myelinating oligodendrocytes don’t form nodal microvilli, allowing glial cells to contact the uninsulated axolemma directly at the nodes of Ranvier. Using immunofluorescent staining and electron microscopy, a recent study revealed direct contact between microglia processes and the nodes in rat corpus callosum^69^, although the physiological role of the contact remains elusive. Here, we assessed microglia-axon contacts at the nodes of Ranvier *in vivo* using Thy1-YFP/Cx3Cr1-GFP double transgenic mice and studied the dynamic behavior of microglia-node interactions during axonal degeneration induced by laser axotomy. Specifically, the position of nodes was first confirmed by merging the SRS image of myelin and the TPEF image of YFP labeled axons. As can be seen, at the nodes of Ranvier the axon is not wrapped by myelin and exhibits a decreased diameter compared with the internode regions (**Fig. 5a**). First, we conducted 1-hour time-lapse multimodal imaging of the spinal cord through the LF window. As expected, the microglial cells under the window displayed ramified morphology with highly motile processes. Interestingly, we observed that a large proportion of nodes (72.4%, n=21) were contacted by microglial processes (**Fig. 5b**). Notably, there were a small number of nodes (17.2%, n=5) showing constant contact with microglia, with one of the microglial processes sticking to the node of Ranvier and remaining stable over time (**Fig. 5c**). Nevertheless, most of the microglia-node contacts were intermittent (55.2%, n=16), occurring as microglial processes randomly scanning over the surrounding environment (**Fig. 5d**). In addition, we also observed that a microglial cell can access two nodes of Ranvier simultaneously with its highly branched processes (**Fig. 5d**), showing the diversity of microglia-node interactions. Then we explored this contact at the nodes of injured axons. Time-lapse imaging was performed for an hour before and almost immediately after laser axotomy on a single axon (**Fig. 5f**). To avoid directly influencing the behavior of microglia around nodes of Ranvier, laser axotomy was performed at least 200 *μ*m distal to the target node (**Supplementary Fig. 17**). After every laser injury, we monitored the dynamics of the microglia around the node and found that their processes were not recruited to the lesion site, suggesting that this precisely controlled distal injury method can exclude the laser-induced microglial response and thus provides an ideal means to study the specific microglial behaviors related to axonal degeneration.

**Figure 5.**
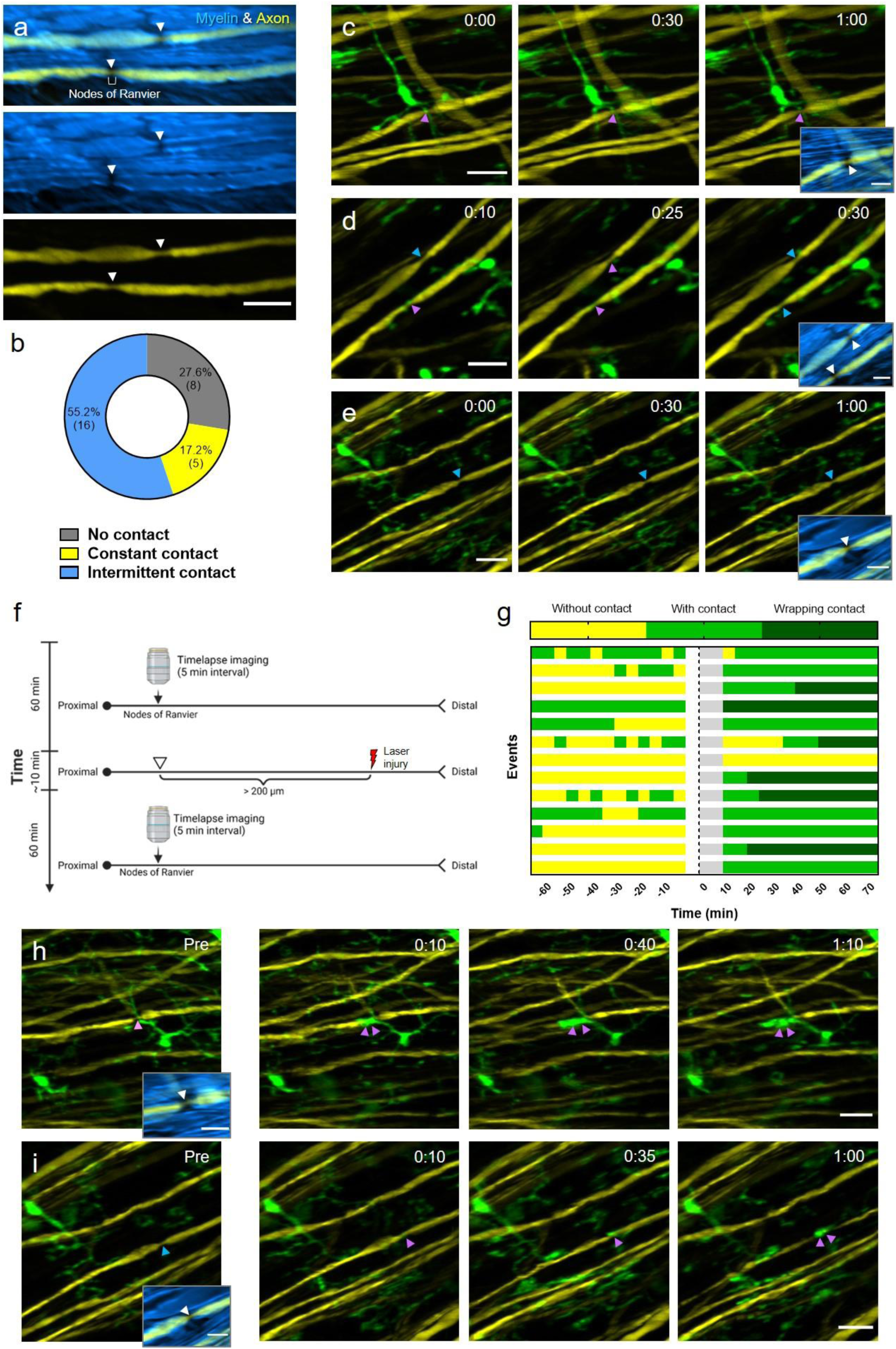
Dynamic contact between microglia and the nodes of Ranvier. (a) The overlay of the SRS image of myelin (blue) and the TPEF image of YFP axon (yellow) shows the structure of the nodes of Ranvier. Arrowheads denote the locations of nodes. Scale bar, 20 *μ*m. (b) The categorization and statistics of microglial contact with the nodes of Ranvier. Totally 29 nodes were studied by 1-hour live imaging. (c-e) Representative results of constant contact (c), intermittent contact (d) and no contact (e) between microglia (green) and the nodes of Ranvier (arrowheads) during 1-hour time-lapse imaging. Insets, overlay of the SRS image of myelin (blue) and the TPEF image of axons (yellow) showing the position of nodes (white arrowheads). Purple arrowheads indicate nodes with microglial contact, while blue arrowheads indicate nodes with no microglial contact. Scale bar, 20 *μ*m. (f) Illustration of the experimental design. Single-axon laser axotomy was conducted at more than 200 *μ*m away from the target node of Ranvier to avoid direct laser-induced microglial activation near the nodes. (g) Quantification of microglia-nodes contact before and after laser axotomy. Each bar represents a node of Ranvier. The blank and grey areas on the bars denote the time for laser injury and imaging setup when imaging was not performed. Laser injury was performed at time 0. (h, i) Representative time sequence images of nodes with wrapped contacts with microglia after axon injury. Before injury, the node in (h) has constant contact with microglia, while the node in (i) has no contact with microglia. After injury, wrapping contacts with nodes enveloped by microglia processes are indicated with double purple arrowheads. Time post injury is presented as hr : min. Scale bar, 20 *μ*m. Insets, overlay of myelin SRS image and axon fluorescence image. Scale bar for all the insets, 10 *μ*m.

Strikingly, we found that after injury at the distal end of the axon, the nodes of Ranvier were contacted by microglial processes significantly more frequently (**Fig. 5g**). Among the 12 nodes of Ranvier which had no contact or intermittent contact with microglia before laser injury, 10 nodes attracted microglial processes within 15 min after axotomy and remained in constant contact during the following hour. Moreover, in some cases, microglial processes were intensively recruited and fused to the nodes, forming an enlarged containment around the nodes (**Fig. 5h,i, supplementary video 3-5**). This wrapping contact was observed in about half the nodes (6/13). These results show that the microglia-node contact is strongly regulated in the injured axons. Although the mechanisms underlying the pronounced changes in microglia-node interactions remain unclear, it is suggested that microglia dynamics can be modulated by the concentration of cations such as potassium (*K*^+^)^32^ and calcium (*Ca*^2+^)^70,71^. At the nodes of Ranvier, where action potentials regenerate, large amount of *K*^+^ and Na^+^ are rapidly exchanged on the uninsulated axolemma^68^. In addition, calcium influx through the nodes is reported to happen in a manner dependent on neuron activity^72,73^. Therefore, microglia contact with nodes of Ranvier may be closely associated with the change of *K*^+^ and/or *Ca*^2+^ concentrations around the nodes. As laser axotomy disrupts the axolemma and surrounding myelin, it would cause a rapid depolarization and an occasional burst of action potentials^74^. Therefore, it is rational to speculate that laser axotomy may affect the concentrations of cations around the node, which further modulates microglia-node contact. Meanwhile, as a unique pathological response triggered by axon injury, the strengthened contact between microglial processes and nodes of Ranvier may offer valuable insights into the regulation of axon-glia interactions during the neurodegeneration process. With this *in vivo* spinal cord imaging method based on the minimal-invasive intervertebral window and multimodal NLO microscopy, we demonstrated a promising way to study the dynamic interaction between the nodes of Ranvier and microglia under normal and injury condition, opening a door for future studies associated with the functions of the nodes of Ranvier.

## 3. DISCUSSION

Since spinal microglia plays a crucial regulatory role in homeostasis^21^, neurodevelopment^75,76^, and neuronal degeneration or regeneration^62,76–78^, during the surgical preparation of the intervertebral window for chronic imaging, it is of great importance to avoid activating the spinal microglia in order to maintain the native microenvironment of the spinal cord. In this study, we demonstrated the use of a minimally invasive intervertebral window with an optical clearing method and NLO microscopy to achieve long-term (167 days), repetitive (16 times), high-resolution (subcellular structure-resolved), and most importantly, inflammation-free (microglia inactive) imaging of mouse spinal cord *in vivo*. To improve the integrity and rigidity of the intervertebral window, we retained the ligamentum flavum to serve as a buffer for any mechanical force to the spinal cord caused by surgery. This is a key procedure to protect the underlying spinal cord tissue and dramatically reduce the possibility of window preparation activating inflammation. A side effect is that newly generated tissues above and below the ligamentum flavum will gradually lower the window’s clarity and reduce the quality of imaging. To solve this problem, we gently removed the newly grown tissues above the ligamentum flavum, and more crucially, we applied an optical clearing method using Iodixanol as the clearing medium to reduce the photon scattering caused by the window and successfully restore subcellular imaging resolution for more than 160 days. Importantly, by monitoring the morphology of microglia after optical clearing using high-resolution two-photon imaging, we confirmed that administering Iodixanol on the surface of the window does not activate an inflammatory response in the spinal cord, making it a reliable way to improve imaging performance without disturbing spinal homeostasis. We also tested widely reported optical clearing agents, glycerol (**Supplementary Fig. 18**) and PEG400 (**Supplementary Fig. 19**). We found that both agents induced activation of microglia and offered limited improvement for two-photon imaging through the window. However, the optical clearing technique based on Iodixanol allows us to remove less tissue from the surface of the window while not compromising imaging quality, thus reducing the risk of activating inflammation by surgery. With these improvements in window preparation, we managed to conduct repetitive surgery without activating microglia with a high success rate (90%, 36/40) and achieved 16 imaging sessions over 167 days, which is sufficient for the longitudinal study of chronic disorders in the spinal cord, such as multiple sclerosis^79,80^, spinal cord injury^81^ and neuropathic pain^82^.

The microglial morphology was used as an *in vivo* indicator of inflammatory activity. It has been reported previously that as well as the two conventional forms of resting and activated state, microglia may display an intermediate state in which cells preserve a branched morphology under pathological stimuli^30^. This is because microglial activation and the resultant morphological transformation is a gradual process, and may have diverse responses to pathological conditions and functional states^31,83^. Indeed, this progressive, heterogeneous alteration in microglial morphology during the activation process may disturb the accuracy of judgments of the activation states of individual microglia based on morphology. Nevertheless, it is widely accepted that inflammation in local tissues can be determined objectively by rigorous statistical analysis of the average morphological parameters of a large population of microglia in the ROI^30,36,47^. Furthermore, it has been observed that microglial activation in response to acute CNS injury is usually rapid and most of the microglial cells near the lesion site can quickly retract processes and even acquire the amoeboid phenotype within a few hours of the stimulus^22,84^. Therefore, in order to assess the extent of inflammatory activation during the preparation of a spinal cord window, we conducted quantitative and statistical characterization of the ramification index and process endpoints of the spinal cord microglia using time-lapse *in vivo* imaging.

By using a home-built multimodal NLO microscope system that integrates TPEF, SHG and SRS imaging, we achieved simultaneous visualization of a variety of structures in and above the spinal cord, including axons, myelin, microglia, blood vessels, collagen, lipid, etc., facilitating our understanding of the remodeling of the complex microenvironment in the intervertebral window during longitudinal imaging. This multimodal imaging plays a crucial role in characterizing the biophysical and biochemical properties of the intervertebral window, monitoring the axon-glia dynamics following laser injury, and identifying the microglial contacts with the nodes of Ranvier. In addition, our two-photon laser microsurgery provides an ideal model for studying spinal cord injury in a well-controlled manner and specifically, single axon injury in the dorsal column area. As well as the advanced imaging tool, another indispensable factor for high-resolution spinal cord imaging is that we established a custom-designed stabilization stage to minimize the influence of mice breathing during imaging, and also applied rigorous image registration algorithms to correct residual motion artifacts.

It should be noted that although the sub-cellular resolution of two-photon spinal cord imaging can be achieved most of the time through tissue removal and optical clearing, clear images may be hard to acquire when newly generated blood vessels in the epidural space are densely distributed right above the ROI. Further, optical clearing showed smaller improvement for SRS imaging compared to two-photon imaging. This probably results from the chromatic aberration introduced by the optical cleared LF window since SRS generation depends critically on the spatial overlap of the pump and Stokes beams at the focal point. To further improve the image quality under the LF window in the future, adaptive optics could be introduced and integrated into the NLO microscope to compensate for the monochromatic and chromatic aberrations caused by the window. It is also worth noting that the effective area of the intervertebral window decreased significantly after 3 months because of the growth of surrounding rigid tissues that are difficult to remove. Therefore, to avoid losing the longitudinal traced ROI due to the decreased FOV, it is preferable to use the central region of the window for extremely long-term imaging. With the future development of advanced microscopy techniques, this proposed optically cleared LF window will serve as a robust and general tool for neuroscientists to understand cellular dynamics in the spinal cord under physiological and pathological conditions in a live mouse model.

## 4. METHODS AND MATERIALS

### Animal preparation

Heterozygous Cx3Cr1-GFP (B6.129P2(Cg)-Cx3cr1tm1Litt/J)^85^ transgenic mice which express EGFP in microglia were used to characterize the inflammatory activation in the spinal cord. To study axon-glia interaction, Cx3Cr1-GFP mice were crossed with Thy1-YFP (Tg(Thy1-YFP)HJrs/J)^86^ mice to generate the Thy1-YFP/Cx3Cr1-GFP transgenic line for simultaneous imaging of axon and microglia in the spinal cord. All the mice used for imaging experiments were 2-6 months old. Before surgery, all required tools were sterilized by autoclaving. All surfaces which would be touched during surgery were disinfected with 70% ethanol. A sterile field was created for surgery by covering the working area of benchtop with sterile drapes. Mice were anesthetized by intraperitoneal (i.p.) injection of ketamine-xylazine mixture (87.5 mg kg^-1^ and 12.5 mg kg^-1^). Hair on the dorsal surface above the thoracic spine was shaved and completely removed using depilating cream. The dorsal surface was disinfected using iodine solution. A small (∼1.5 cm) midline incision of the skin was made over the T11-T13 vertebra to expose the dorsal tissue (**Supplementary Fig. 4b**). Muscles and tendons on both the top and sides were severed so that the spine can be held stably by clamping the vertebra with two stainless steel clamping bars on a custom-designed stabilization stage (**Supplementary Fig. 4c**). During the surgery, sterile gauze pads and sterile saline were used to control bleeding and clean the wound. The surface of the stabilization stage was maintained at around 37° through a heating pad to keep mice warm during surgery. All animal procedures performed in this work were conducted according to the guidelines of the Laboratory Animal Facility of the Hong Kong University of Science and Technology (HKUST) and were approved by the Animal Ethics Committee of HKUST.

#### Intervertebral window

The surgical procedures for preparing conventional intervertebral windows were modified according to a previous protocol^19,20^. Briefly, the muscle tissues and tendons in the cleft between the vertebra arcs T12 and T13 were completely removed. The ligamentum flavum was carefully peeled using a fine-tip tweezer, while the dura was left intact. The exposed spinal cord was kept moist by irrigating with saline. To prepare the improved intervertebral window with ligamentum flavum, care should be taken to keep the ligamentum flavum unblemished when removing the muscle and tendon in the intervertebral space (**Supplementary Fig. 4d**). In particular, after the window with ligamentum flavum has been exposed, the tweezer tip should not touch the surface of the window during surgery. This is important to avoid inducing microglia activation. Moreover, when cleaning tissue with a saline flush and gauze pad, direct contact with the window surface should also be avoided. To prepare for the imaging, a coverslip was then placed on the clamping bar, and the interspace between the coverslip and the spinal cord was filled with saline or Iodixanol (**Supplementary Fig. 4e**). After imaging, the medium below the coverslip was removed and the surgical area was carefully cleaned using saline and gauze pads. The top area of the surgical window was then covered by liquid Kwik-Sil (World Precision Instruments) to protect it (**Supplementary Fig. 4h**). After the Kwik-Sil got cured (∼3min), the skin on the surgical site was sutured and covered with burn cream (Betadine) to protect from infection. Mice were placed on a heating pad until they recovered fully from anesthesia. For reimaging through the same intervertebral window with ligamentum flavum, the sutured skin was reopened and the covering Kwik-Sil gel was removed. Tissues adhering to the side of the T11-T13 vertebra were detached to enable stable clamping of the spine. If reimaging was performed within four days of the initial surgery, granulation tissue had not formed at the surgical site. Therefore, we only need to clean the window surface by flushing saline and remove loose tissue debris from the surface. With the growth of granulation tissue accompanied by angiogenesis and fibroplasia, the tissues on the surface of the surgical site should be peeled off to expose the ligamentum flavum, which can be easily distinguished from the newly generated tissues by its tough collagenous structures. In addition, the laminae and processes of two vertebrae around the window should always be scraped clean without tissue adhesions. The procedures for imaging and post-imaging preparations are the same as previously described.

#### Dorsal column crush

The spinal cord dorsal column crush was conducted following previous protocols^62,87^ with slight modifications. Briefly, T12 laminectomy was performed to expose the spinal cord using Dumont #2 Laminectomy forceps. Two small holes were made in the dura with a 30-gauge needle symmetrically around 0.5 mm lateral to the midline. A dorsal hemi-crush injury was made by inserting the modified Dumont #5 forcep through the two small holes approximately 0.6 mm into the dorsal spinal cord and squeezing with pressure for 5s, and repeating three times.

### Multimodal NLO microscopy

The setup of our multimodal NLO microscope is shown in **Supplementary Fig. 1**. An integrated optical parametric oscillator (OPO, picoEmerald S, APE) was used as the light source for SRS imaging. It consists of a Stokes beam (1031nm) and pump beam (tunable from 780nm to 960nm) with 2 ps pulse duration and 80 MHz repetition rate. The intensity of the Stokes beam was modulated at 20 MHz by a built-in electro optical modulator. The pump beam was combined with the Stokes beam using a dichroic mirror (D1) inside the picoEmerald S. A femtosecond Ti:sapphire laser (Chameleon Ultra II, Coherent) tuned to 920nm was used as the laser source for exciting two-photon fluorescence and generating second harmonic generation signals. The fs beam was rotated from horizontal to vertical polarization by a half-wave plate(SAHWP05M-1700, Thorlabs) and then combined with the ps beam by a polarizing beam splitter (CCM1-PBS252/M, Thorlabs). The ps beam and fs beam were collimated and magnified by a pair of achromatic doublets to match the 3 mm Galvo XY-scan mirror (6215H, Cambridge Technology). The Galvo mirror and the rear pupil of the objective lens (XLPLN25XSVMP2, 25×/1.05 NA, Olympus) were conjugated by a telecentric scan lens L5 (SL50-CLS2, Thorlabs) and an infinity-corrected tube lens L6 (TTL200-S8, Thorlabs). The laser beam was expanded by the scan and tube lens to fill the back aperture of the objective.

For SRS imaging, the backscattered pump beam collected by the objective was reflected by a polarizing beam splitter (CCM1-PBS252/M, Thorlabs) and directed to a large area (10mm×10mm) Si photodiode (S3590-08; Hamamatsu). A dichroic short-pass filter D3 (69-206, short-pass at 700nm, Edmund) was used to separate the SRS detection path from the fluorescence detection path. A filter set (Fs1) including a short-pass filter (86-108, short-pass at 975nm OD4, Edmund) and a band-pass filter (FF01-850/310, Semrock) were placed before the photodiode to completely block the Stokes beam. The output of the photodiode was then fed into a lock-in amplifier (LIA) for signal demodulation and amplification to obtain highly sensitive detection of stimulated Raman loss (SRL).

For two-photon imaging, the polarizing beam splitter above the objective was replaced by a dichroic beam splitter D2 (FF665-Di02, Semrock) to reflect the TPEF and SHG signal to the photodetection unit. An interchangeable dichroic beam splitter D4 (FF488-Di01-25×36 or FF518-Di01-25×36, Semrock) was placed after D3 to separate the fluorescence into two current photomultiplier (PMT) modules (H11461-03 and H11461-01, Hamamatsu). Two filter sets Fs2 (FF01-715/SP-25, Semrock; FF01-525/50, Semrock or HQ620/60X, Chroma) and Fs3 (FF01-720/SP-25, Semrock; FF01-525/50, Semrock or HQ440/80M, Chroma) were placed before the PMTs to reject the excitation beam and transmit fluorescence. The output currents of the two PMTs were then converted to voltage by two current amplifiers (SR570, Stanford research). The outputs of the two current amplifiers and LIA were then fed into a multifunction acquisition card (PCIe-6363, National Instrument) to reconstruct the image. For spectral characterization of emitted TPEF, the dichroic mirror D4 was switched to 665dcxr (Chroma) to reflect fluorescence onto a fiber-based spectroscopic detection module. The reflected fluorescence was filtered by a short pass filter (SP01-633RU-25, Semrock) and coupled into a fiber bundle before being directed to a multispectral detection system consisting of a spectrograph (455 ∼ 650 nm) equipped with a 16-channel PMT module (PML-16-C-0, Becker & Hickl). This detection system allows spectral measurements for each pixel of the TPEF image with a 13-nm spectral resolution. All the hardware was controlled by a custom-written C# program to acquire two-photon and SRS images.

The hyperspectral SRS sweeping mode was used to acquire the SRS spectra of solutions and tissues in the fingerprint and C-H stretch region. First, temporal overlapping calibration of the pump and Stokes beams at the fingerprint and C-H vibration regions was performed by adjusting a built-in delay stage based on the SRS signal of 6 μm polystyrene beads (Polysciences, Inc., Warrington, PA), Olive oil and heavy water (99% pure, D2O) at their specific Raman peaks. Since solution samples are homogenous with little scattering, to achieve SRS imaging of solutions in an epi-detection configuration, a piece of folded tissue paper was stuck to the bottom of the slide to backscatter the SRS signals. The wavelength of the pump beam was sequentially tuned with 0.3-nm steps by the program through a serial communication port. For Iodixanol SRS imaging, the Lyot filter inside the laser was adjusted to fast tune the pump wavelength from 891.1 nm (1523 cm^-1^, on-resonant) to 893 nm (1500 cm^-1^, off-resonant). By synchronizing the Lyot filter to the frame trigger, a pair of “on-resonant” and “off-resonant” images could be acquired with less than 3 s switching time. The final Iodixanol image was obtained by subtracting the off-resonant signals from the on-resonant signals.

### *In vivo* imaging

Before each imaging session, the mouse received a retro-orbital intravenous injection of 100 μl Texas Red dextran (70 kDa, 1mg/100ul in saline, Invitrogen) to label blood vessels when necessary. The stabilization stage securing the mouse was placed on a five-axis stage beneath our customized microscope. The five-axis stage allows three-axis translation and ±5° pitch and roll flexure motion. To reduce the motion artifacts caused by breathing, the mouse’s head was secured by two head bars and the mouse’s body was elevated a little by lowering the holding plate to allow room for chest movement during breathing (**Supplementary Fig. 4a**). The mouse’s spinal cord was aligned perpendicular to the objective axis by adjusting the roll and pitch angles of the stage guided by the bright-field image (4× objective, 0.16 NA, Olympus). Since the spinal cord has a natural curvature, in order to align the sample surface precisely during NLO imaging over a large FOV, the angle needs to be finely adjusted for each small sub-region guided by the TPEF signal of each FOV. The femtosecond laser was tuned to 920 nm for TPEF excitation of GFP, YFP or Texas Red. First, a 10× objective lens (NA = 0.45, Nikon) was used to obtain an image of the entire intervertebral window as a roadmap for navigating between imaging sessions. Then a 25× water immersion objective (NA = 1.05, Olympus) was used for high-resolution two-photon and SRS imaging of the target area. For two-photon imaging with 10× and 25× objectives, the post-objective power ranged from 40 to 65 mw and 10 to 50 mw respectively, depending on the clarity of the intervertebral window. During imaging, the holding plate was heated to 37°C to keep the mouse warm. Ketamine-xylazine (43.75 mg kg^-1^; 6.25 mg kg^-1^)) were supplemented when necessary.

### Optical clearing by Iodixanol

Iodixanol/OptiPrep (D1556, Sigma-Aldrich) was purchased as a 60% w/v stock solution and prepared with various concentrations by diluting the 60% w/v stock solution in sterile phosphate buffered saline (PBS). To achieve optical clearing of the IWLF, Iodixanol was applied and supplemented hourly. Imaging was usually started 10 min after every Iodixanol administration when the optical clearing effect reached a plateau.

### Histology

Mice were deeply anesthetized and then perfused transcardially with 20 ml PBS to wash out the blood and 20 ml 4% (w/v) PFA (Sigma-Aldrich) for fixation. Spinal cord segments (∼1 cm) at the surgical and control region were dissected out and immersed in 15% (w/v) sucrose PBS solution for 12 hrs before further dehydration in 30% sucrose PBS solution. After sedimentation, samples were frozen at −80°C and then cut to 50 um-thick sagittal sections on a CryoStar NX70 Cryostat (Thermo Scientific). The GFP-labeled microglia cells located less than 50 *μ*m below the dorsal surface were imaged for morphological analysis.

### Laser axotomy

Laser axotomy is achieved by a highly localized nonlinear process based on multi-photon ionization and plasma-mediated ablation^88^. To perform imaging-guided laser axotomy, a femtosecond laser tuned to 920 nm was focused on the targeted axon for 1-4 s with an average power of 250 mW. The lesion caused by laser ablation can be visualized and quantified by the newly generated fluorescence^88,89^ or SRS signal of the spinal cord.

### Image processing and analysis

Since *in vivo* spinal cord imaging would be affected severely by motion artifacts caused by breathing and heartbeats, it is necessary to perform image registration to acquire stable images. For multimodal NLO imaging, three-dimensional (3D) optical sectioning was performed to obtain images at different depths. To reduce intra-frame distortion, 10 frames (512×512 pixels) were acquired per slice with a 2-Hz frame rate. Single-channel 3D image registration was carried out as follows. First, image registration was performed on the sequential frames for each slice using the ‘StackReg’ plugin^90^ in Fiji software^91^. Then the registered frames were averaged to form a target image used for the next step of registration. Using the ‘bUnwarpJ’ plugin^92^ in Fiji, each raw image frame was then registered individually to the target image. The registered frames were then averaged to obtain the final slice for each depth with minimal motion artifacts and improved signal-to-background ratio. Registration between slices was then performed using the ‘StackReg’ plugin to construct the 3D image stack. For two-channel 3D image registration, since each pair of the two-channel images was acquired simultaneously with the same deformation, they should be registered using the same transformation parameters. During ‘bUnwarpJ’ registration, the transformation information for each frame of the single-channel images was recorded and then applied to the corresponding frames of another channel. To co-localize SRS image and fs laser excited TPEF image, we captured the ps laser excited TPEF images simultaneously with the SRS image, which were then used as reference images for SRS and fs TPEF image registration. The ‘MultiStackReg’ plugin in Fiji was used for multichannel stack registration using the same transformation parameters. All these registration procedures were implemented in the Fiji macro programming language. To acquire the large-FOV images of the intervertebral window as shown in **Fig.1c, Fig. 4a, Supplementary Fig. 4a, Supplementary Fig. 15a, Supplementary Fig. 16** and **Supplementary Fig. 17**, the sub-images were stitched using the ‘Pairwise stitching’ plugin^93^ in Fiji.

To characterize the image contrast, fluorescence images were stacked as a maximum z projection. In each FOV, microglia with visible cell bodies were randomly selected for contrast characterization. An intensity profile was plotted through the center of the microglia cell body. The peak value of the profile is represented as *I*_0_. The background value B is defined as the averaged intensity value over a region at 10 *μ*m away from the intensity peak^94^. The image contrast can be calculated as^16^:

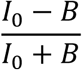

A contrast value of 0 represents no contrast while 1 represents noiseless contrast.

For quantification of microglia ramification index and process endpoints, microglia with intact and clear morphology (contrast ≥ _0_.97) (**Supplementary Fig 13**) in each FOV were randomly selected for morphological quantification without bias. The microglial ramification index and process endpoints were quantified based on the published methods^32,33^ with small modifications. Briefly, binary images of individual microglia were first acquired and saved as independent files as previously described^32^. Microglial binary images were skeletonized using the ‘bwskel’ MATLAB function and the endpoints of each microglial skeleton were counted using the ‘bwmorph’ MATLAB function. To quantify the ramification index, the MATLAB functions ‘bwarea’ and ‘bwperim’ (8-connected neighborhood) were used to acquire the area and perimeter value of each cell. The ramification index is then calculated as^32^:

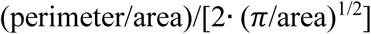

For analysis of microglia aggregation and axonal degeneration, a dot rectangular region of interest was outlined along the axonal degeneration path to quantify the spatial distribution of the microglial aggregation after laser axotomy (**Fig. 4a**). The maximal z projection images of axon and microglia in the rectangle area were smoothed and converted to a bit-map. The aggregation size of the microglia was calculated as the sum of the GFP fluorescence signals in the bit-mapped area normalized by the fluorescence intensity of surrounding uninjured axons. The axonal diameter was measured based on the YFP fluorescence signals along the degeneration path.

For microglia-nodes of Ranvier contact analysis, the positions of the nodes of Ranvier on the axon were first confirmed by merging the myelin SRS image and axon TPEF image. Axon images were then merged with microglia images to visualize microglia contacts with the nodes of Ranvier. To reduce noise, images were smoothed after background subtraction. Contacts were defined as the 3D colocalization of the fluorescence of microglia and the node of Ranvier. Wrapping contact was identified when the nodes of Ranvier were totally enveloped by microglia processes.

### Statistical analysis

Statistical analysis and data visualization were performed using GraphPad Prism 7 software. All the data are presented as mean ± s.e.m. and *α* =0.05 for all analyses. P values for ordinary one-way ANOVA with Dunnett’s multiple comparison test (for normally distributed data) or Kruskal-Wallis test (for non-normally distributed data) are given on the figures. Data normality was checked using Shapiro-Wilk normality test. No statistical methods were used to predetermine sample sizes, but our sample sizes were similar to reported studies of chronic spinal cord imaging^16,17^.

## Acknowledgments

This work was supported by the Hong Kong Research Grants Council through grants 16103215, 16148816, 16102518, 16102920, T13-607/12R, T13-706/11-1, T13-605/18W, C6002-17GF, C6001-19E, N_HKUST603/19 and the Innovation and Technology Commission (ITCPD/17-9), and the Area of Excellence Scheme of the University Grants Committee (AoE/M-604/16, AOE/M-09/12) and the Hong Kong University of Science & Technology (HKUST) through grant RPC10EG33.

## Author contributions

W.W, S.H., and J.Y.Q conceived the research idea and designed the experiments; X.L. and W.W. built the imaging systems; W.W performed animal surgery and imaging experiments; J.W. perfused the animal and prepared slices for histology analysis; W.W analyzed the data with the help of C.C; S.H. and W.W. wrote the paper with input from all other authors.

## REFERENCE

1. Kobat, D. et al. Deep tissue multiphoton microscopy using longer wavelength excitation. Opt. Express, OE 17, 13354–13364 (2009).

2. Xu, H.-T., Pan, F., Yang, G. & Gan, W.-B. Choice of cranial window type for in vivo imaging affects dendritic spine turnover in the cortex. Nature Neuroscience 10, 549–551 (2007).

3. Sohler, T. P., Lothrop, G. N. & Forbes, H. S. The Pial Circulation of Normal, Non-Anesthetized Animals Part Ii. the Effects of Drugs, Alcohol and Co2. J Pharmacol Exp Ther 71, 331–335 (1941).

4. Drew, P. J. et al. Chronic optical access through a polished and reinforced thinned skull. Nature Methods 7, 981–984 (2010).

5. Yang, G., Pan, F., Parkhurst, C. N., Grutzendler, J. & Gan, W.-B. Thinned-skull cranial window technique for long-term imaging of the cortex in live mice. Nat Protoc 5, 201–208 (2010).

6. Grutzendler, J., Kasthuri, N. & Gan, W.-B. Long-term dendritic spine stability in the adult cortex. Nature 420, 812–816 (2002).

7. Li, Y., Du, X. & Du, J. Resting microglia respond to and regulate neuronal activity in vivo. Communicative & Integrative Biology 6, e24493 (2013).

8. Hamm, J. P., Peterka, D. S., Gogos, J. A. & Yuste, R. Altered Cortical Ensembles in Mouse Models of Schizophrenia. Neuron 94, 153-167.e8 (2017).

9. Xu, Z. et al. Rescue of maternal immune activation-induced behavioral abnormalities in adult mouse offspring by pathogen-activated maternal T reg cells. Nature Neuroscience 1–13 (2021) doi:10.1038/s41593-021-00837-1.

10. Misgeld, T., Nikic, I. & Kerschensteiner, M. In vivo imaging of single axons in the mouse spinal cord. Nat Protoc 2, 263–268 (2007).

11. Davalos, D. et al. Stable in vivo imaging of densely populated glia, axons and blood vessels in the mouse spinal cord using two-photon microscopy. Journal of Neuroscience Methods 169, 1–7 (2008).

12. Ran, C., Hoon, M. A. & Chen, X. The coding of cutaneous temperature in the spinal cord. Nature Neuroscience 19, 1201–1209 (2016).

13. Kerschensteiner, M., Schwab, M. E., Lichtman, J. W. & Misgeld, T. In vivo imaging of axonal degeneration and regeneration in the injured spinal cord. Nature Medicine 11, 572–577 (2005).

14. Dray, C., Rougon, G. & Debarbieux, F. Quantitative analysis by in vivo imaging of the dynamics of vascular and axonal networks in injured mouse spinal cord. Proceedings of the National Academy of Sciences 106, 9459–9464 (2009).

15. Di Maio, A. et al. In Vivo Imaging of Dorsal Root Regeneration: Rapid Immobilization and Presynaptic Differentiation at the CNS/PNS Border. Journal of Neuroscience 31, 4569–4582 (2011).

16. Farrar, M. J. et al. Chronic in vivo imaging in the mouse spinal cord using an implanted chamber. Nature Methods 9, 297–302 (2012).

17. Fenrich, K. K. et al. Long-term in vivo imaging of normal and pathological mouse spinal cord with subcellular resolution using implanted glass windows. The Journal of Physiology 590, 3665–3675 (2012).

18. Figley, S. A. et al. A Spinal Cord Window Chamber Model for In Vivo Longitudinal Multimodal Optical and Acoustic Imaging in a Murine Model. PLOS ONE 8, e58081 (2013).

19. Kim, J. V. et al. Two-photon laser scanning microscopy imaging of intact spinal cord and cerebral cortex reveals requirement for CXCR6 and neuroinflammation in immune cell infiltration of cortical injury sites. Journal of Immunological Methods 352, 89–100 (2010).

20. Nadrigny, F., Le Meur, K., Schomburg, E. D., Safavi-Abbasi, S. & Dibaj, P. Two-Photon Laser-Scanning Microscopy for Single and Repetitive Imaging of Dorsal and Lateral Spinal White Matter In Vivo. Physiol Res 531–537 (2017) doi:10.33549/physiolres.933461.

21. Nimmerjahn, A., Kirchhoff, F. & Helmchen, F. Resting Microglial Cells Are Highly Dynamic Surveillants of Brain Parenchyma in Vivo. Science 308, 1314–1318 (2005).

22. Davalos, D. et al. ATP mediates rapid microglial response to local brain injury in vivo. Nature Neuroscience 8, 752–758 (2005).

23. Dibaj, P. et al. NO mediates microglial response to acute spinal cord injury under ATP control in vivo. Glia 58, 1133–1144 (2010).

24. Stollg, G. & Jander, S. The role of microglia and macrophages in the pathophysiology of the CNS. Progress in Neurobiology 58, 233–247 (1999).

25. Perry, V. H., Nicoll, J. A. R. & Holmes, C. Microglia in neurodegenerative disease. Nat Rev Neurol 6, 193–201 (2010).

26. Kreutzberg, G. W. Microglia: a sensor for pathological events in the CNS. Trends in Neurosciences 19, 312–318 (1996).

27. Stence, N., Waite, M. & Dailey, M. E. Dynamics of microglial activation: A confocal time-lapse analysis in hippocampal slices. Glia 33, 256–266 (2001).

28. Davis, E. J., Foster, T. D. & Thomas, W. E. Cellular forms and functions of brain microglia. Brain Research Bulletin 34, 73–78 (1994).

29. Karperien, A., Ahammer, H. & Jelinek, H. Quantitating the subtleties of microglial morphology with fractal analysis. Front. Cell. Neurosci. 7, (2013).

30. Fernández-Arjona, M. del M., Grondona, J. M., Fernández-Llebrez, P. & López-Ávalos, M. D. Microglial Morphometric Parameters Correlate With the Expression Level of IL-1β, and Allow Identifying Different Activated Morphotypes. Front. Cell. Neurosci. 13, (2019).

31. Gomez-Nicola, D. & Perry, V. H. Microglial Dynamics and Role in the Healthy and Diseased Brain: A Paradigm of Functional Plasticity. Neuroscientist 21, 169–184 (2015).

32. Madry, C. et al. Microglial Ramification, Surveillance, and Interleukin-1β Release Are Regulated by the Two-Pore Domain K+ Channel THIK-1. Neuron 97, 299-312.e6 (2018).

33. Morrison, H. W. & Filosa, J. A. A quantitative spatiotemporal analysis of microglia morphology during ischemic stroke and reperfusion. J Neuroinflammation 10, 4 (2013).

34. Stowell, R. D. et al. Noradrenergic signaling in the wakeful state inhibits microglial surveillance and synaptic plasticity in the mouse visual cortex. Nature Neuroscience 22, 1782–1792 (2019).

35. Liu, Y. U. et al. Neuronal network activity controls microglial process surveillance in awake mice via norepinephrine signaling. Nat Neurosci 22, 1771–1781 (2019).

36. Sun, W. et al. In vivo Two-Photon Imaging of Anesthesia-Specific Alterations in Microglial Surveillance and Photodamage-Directed Motility in Mouse Cortex. Front. Neurosci. 13, (2019).

37. Sołtys, Z., Ziaja, M., Pawliński, R., Setkowicz, Z. & Janeczko, K. Morphology of reactive microglia in the injured cerebral cortex. Fractal analysis and complementary quantitative methods. Journal of Neuroscience Research 63, 90–97 (2001).

38. Zanier, E. R., Fumagalli, S., Perego, C., Pischiutta, F. & De Simoni, M.-G. Shape descriptors of the “never resting” microglia in three different acute brain injury models in mice. Intensive Care Med Exp 3, (2015).

39. Soltys, Z. et al. Quantitative morphological study of microglial cells in the ischemic rat brain using principal component analysis. Journal of Neuroscience Methods 146, 50–60 (2005).

40. Sousa, A. A. de et al. Three-dimensional morphometric analysis of microglial changes in a mouse model of virus encephalitis: age and environmental influences. European Journal of Neuroscience 42, 2036–2050 (2015).

41. Riester, K. et al. In vivo characterization of functional states of cortical microglia during peripheral inflammation. Brain, Behavior, and Immunity (2019) doi:10.1016/j.bbi.2019.12.007.

42. Norden, D. M., Trojanowski, P. J., Villanueva, E., Navarro, E. & Godbout, J. P. Sequential activation of microglia and astrocyte cytokine expression precedes increased iba-1 or GFAP immunoreactivity following systemic immune challenge. Glia 64, 300–316 (2016).

43. Kozlowski, C. & Weimer, R. M. An Automated Method to Quantify Microglia Morphology and Application to Monitor Activation State Longitudinally In Vivo. PLoS One 7, (2012).

44. Staikopoulos, V. et al. Graded peripheral nerve injury creates mechanical allodynia proportional to the progression and severity of microglial activity within the spinal cord of male mice. Brain, Behavior, and Immunity S0889159120323941 (2020) doi:10.1016/j.bbi.2020.11.018.

45. Hamilton, N. et al. The failure of microglia to digest developmental apoptotic cells contributes to the pathology of RNASET2-deficient leukoencephalopathy. Glia 68, 1531–1545 (2020).

46. Neubrand, V. E., Forte-Lago, I., Caro, M. & Delgado, M. The atypical RhoGTPase RhoE/Rnd3 is a key molecule to acquire a neuroprotective phenotype in microglia. Journal of Neuroinflammation 15, 343 (2018).

47. Heindl, S. et al. Automated Morphological Analysis of Microglia After Stroke. Front. Cell. Neurosci. 12, (2018).

48. Lafrenaye, A. D., Todani, M., Walker, S. A. & Povlishock, J. T. Microglia processes associate with diffusely injured axons following mild traumatic brain injury in the micro pig. Journal of Neuroinflammation 12, 186 (2015).

49. Takatsuru, Y., Nabekura, J., Ishikawa, T., Kohsaka, S. & Koibuchi, N. Early-life stress increases the motility of microglia in adulthood. J Physiol Sci 65, 187–194 (2015).

50. Morrison, H., Young, K., Qureshi, M., Rowe, R. K. & Lifshitz, J. Quantitative microglia analyses reveal diverse morphologic responses in the rat cortex after diffuse brain injury. Sci Rep 7, 13211 (2017).

51. Olszewski, A. D., Yaszemski, M. J. & White, A. A. I. The Anatomy of the Human Lumbar Ligamentum Flavum: New Observations and Their Surgical Importance. Spine 21, 2307–2312 (1996).

52. Saito, T. et al. Experimental Mouse Model of Lumbar Ligamentum Flavum Hypertrophy. PLOS ONE 12, e0169717 (2017).

53. Kreisel, D. et al. In vivo two-photon imaging reveals monocyte-dependent neutrophil extravasation during pulmonary inflammation. Proceedings of the National Academy of Sciences 107, 18073–18078 (2010).

54. Boothe, T. et al. A tunable refractive index matching medium for live imaging cells, tissues and model organisms. Elife 6, (2017).

55. Priebe, H. et al. Synthesis and Characterization of Iodixanol. Acta Radiol 36, 21–31 (1995).

56. Li, X. et al. Quantitative Imaging of Lipid Synthesis and Lipolysis Dynamics in Caenorhabditis elegans by Stimulated Raman Scattering Microscopy. Analytical Chemistry 91, 2279–2287 (2019).

57. Li, X., Jiang, M., Lam, J. W. Y., Tang, B. Z. & Qu, J. Y. Mitochondrial Imaging with Combined Fluorescence and Stimulated Raman Scattering Microscopy Using a Probe of the Aggregation-Induced Emission Characteristic. J. Am. Chem. Soc. 139, 17022–17030 (2017).

58. Albrechtsson, U., Lärusdóttir, H., Norgren, L. & Lundby, B. Iodixanol — a New Nonionic Dimer — in Aortofemoral Angiography. Acta Radiologica 33, 611–613 (1992).

59. Heglund, I. F., Michelet, Å. A., Blazak, W. F., Furuhama, K. & Holtz, E. Preclinical Pharmacokinetics and General Toxicology of Iodixanol. Acta Radiol 36, 69–82 (1995).

60. Ford, T., Graham, J. & Rickwood, D. Iodixanol: A Nonionic Iso-osmotic Centrifugation Medium for the Formation of Self-Generated Gradients. Analytical Biochemistry 220, 360–366 (1994).

61. Chen, C. et al. High-resolution two-photon transcranial imaging of brain using direct wavefront sensing. Photon. Res. 9, 1144 (2021).

62. Evans, T. A. et al. High-resolution intravital imaging reveals that blood-derived macrophages but not resident microglia facilitate secondary axonal dieback in traumatic spinal cord injury. Experimental Neurology 254, 109–120 (2014).

63. Horn, K. P., Busch, S. A., Hawthorne, A. L., van Rooijen, N. & Silver, J. Another barrier to regeneration in the CNS: Activated macrophages induce extensive retraction of dystrophic axons through direct physical interactions. J Neurosci 28, 9330–9341 (2008).

64. Wang, X. et al. Macrophages in spinal cord injury: Phenotypic and functional change from exposure to myelin debris. Glia 63, 635–651 (2015).

65. Greenhalgh, A. D. & David, S. Differences in the Phagocytic Response of Microglia and Peripheral Macrophages after Spinal Cord Injury and Its Effects on Cell Death. J. Neurosci. 34, 6316–6322 (2014).

66. Busch, S. A., Horn, K. P., Silver, D. J. & Silver, J. Overcoming Macrophage-Mediated Axonal Dieback Following CNS Injury. J. Neurosci. 29, 9967–9976 (2009).

67. Landon, D. N. & Williams, P. L. Ultrastructure of the Node of Ranvier. Nature 199, 575–577 (1963).

68. Lubetzki, C., Sol-Foulon, N. & Desmazières, A. Nodes of Ranvier during development and repair in the CNS. Nat Rev Neurol 16, 426–439 (2020).

69. Zhang, J., Yang, X., Zhou, Y., Fox, H. & Xiong, H. Direct contacts of microglia on myelin sheath and Ranvier’s node in the corpus callosum in rats. J Biomed Res 33, 192–200 (2019).

70. Eyo, U. B. et al. Modulation of Microglial Process Convergence Toward Neuronal Dendrites by Extracellular Calcium. J. Neurosci. 35, 2417–2422 (2015).

71. Chattopadhyay, N. et al. The Extracellular Calcium-Sensing Receptor Is Expressed in Rat Microglia and Modulates an Outward K+ Channel. Journal of Neurochemistry 72, 1915–1922 (1999).

72. Gründemann, J. & Clark, B. A. Calcium-Activated Potassium Channels at Nodes of Ranvier Secure Axonal Spike Propagation. Cell Reports 12, 1715–1722 (2015).

73. Zhang, Z. & David, G. Stimulation-induced Ca2+ influx at nodes of Ranvier in mouse peripheral motor axons. The Journal of Physiology 594, 39–57 (2016).

74. Berdan, R. C., Easaw, J. C. & Wang, R. Alterations in membrane potential after axotomy at different distances from the soma of an identified neuron and the effect of depolarization on neurite outgrowth and calcium channel expression. Journal of Neurophysiology 69, 151–164 (1993).

75. Cserép, C. et al. Microglia monitor and protect neuronal function through specialized somatic purinergic junctions. Science 367, 528–537 (2020).

76. Cserép, C., Pósfai, B. & Dénes, Á. Shaping Neuronal Fate: Functional Heterogeneity of Direct Microglia-Neuron Interactions. Neuron 109, 222–240 (2021).

77. Neumann, H., Kotter, M. R. & Franklin, R. J. M. Debris clearance by microglia: an essential link between degeneration and regeneration. Brain 132, 288–295 (2009).

78. Lloyd, A. F. et al. Central nervous system regeneration is driven by microglia necroptosis and repopulation. Nat Neurosci 22, 1046–1052 (2019).

79. Davalos, D. et al. Fibrinogen-induced perivascular microglial clustering is required for the development of axonal damage in neuroinflammation. Nature Communications 3, 1–15 (2012).

80. Bartholomäus, I. et al. Effector T cell interactions with meningeal vascular structures in nascent autoimmune CNS lesions. Nature 462, 94–98 (2009).

81. Park, K. K. et al. Promoting Axon Regeneration in the Adult CNS by Modulation of the PTEN/mTOR Pathway. Science 322, 963–966 (2008).

82. Chen, G., Zhang, Y.-Q., Qadri, Y. J., Serhan, C. N. & Ji, R.-R. Microglia in Pain: Detrimental and Protective Roles in Pathogenesis and Resolution of Pain. Neuron 100, 1292–1311 (2018).

83. Petersen, M. A. & Dailey, M. E. Diverse microglial motility behaviors during clearance of dead cells in hippocampal slices. Glia 46, 195–206 (2004).

84. Kawabori, M. & Yenari, M. A. The role of the microglia in acute CNS injury. Metab Brain Dis 30, 381–392 (2015).

85. Jung, S. et al. Analysis of Fractalkine Receptor CX3CR1 Function by Targeted Deletion and Green Fluorescent Protein Reporter Gene Insertion. Molecular and Cellular Biology 20, 4106–4114 (2000).

86. Feng, G. et al. Imaging Neuronal Subsets in Transgenic Mice Expressing Multiple Spectral Variants of GFP. Neuron 28, 41–51 (2000).

87. Chen, W. et al. Rapamycin-Resistant mTOR Activity Is Required for Sensory Axon Regeneration Induced by a Conditioning Lesion. eneuro 3, ENEURO.0358-16.2016 (2016).

88. Qin, Z. et al. New fluorescent compounds produced by femtosecond laser surgery in biological tissues: the mechanisms. Biomed. Opt. Express, BOE 9, 3373–3390 (2018).

89. Sun, Q. et al. In vivo imaging-guided microsurgery based on femtosecond laser produced new fluorescent compounds in biological tissues. Biomedical Optics Express 9, 581 (2018).

90. Thevenaz, P., Ruttimann, U. E. & Unser, M. A pyramid approach to subpixel registration based on intensity. IEEE Transactions on Image Processing 7, 27–41 (1998).

91. Schindelin, J. et al. Fiji: an open-source platform for biological-image analysis. Nat Methods 9, 676–682 (2012).

92. Arganda-Carreras, I. et al. Consistent and Elastic Registration of Histological Sections Using Vector-Spline Regularization.in Computer Vision Approaches to Medical Image Analysis (eds. Beichel, R. R. & Sonka, M.) vol. 4241 85–95 Springer Berlin Heidelberg, 2006).

93. Preibisch, S., Saalfeld, S. & Tomancak, P. Globally optimal stitching of tiled 3D microscopic image acquisitions. Bioinformatics 25, 1463–1465 (2009).

94. Kobat, D., Horton, N. G. & Xu, C. In vivo two-photon microscopy to 1.6-mm depth in mouse cortex. J. Biomed. Opt. 16, 106014 (2011).

