## Supplementary Figures for "Long-term *in vivo* imaging of mouse spinal cord through an optically cleared intervertebral window"

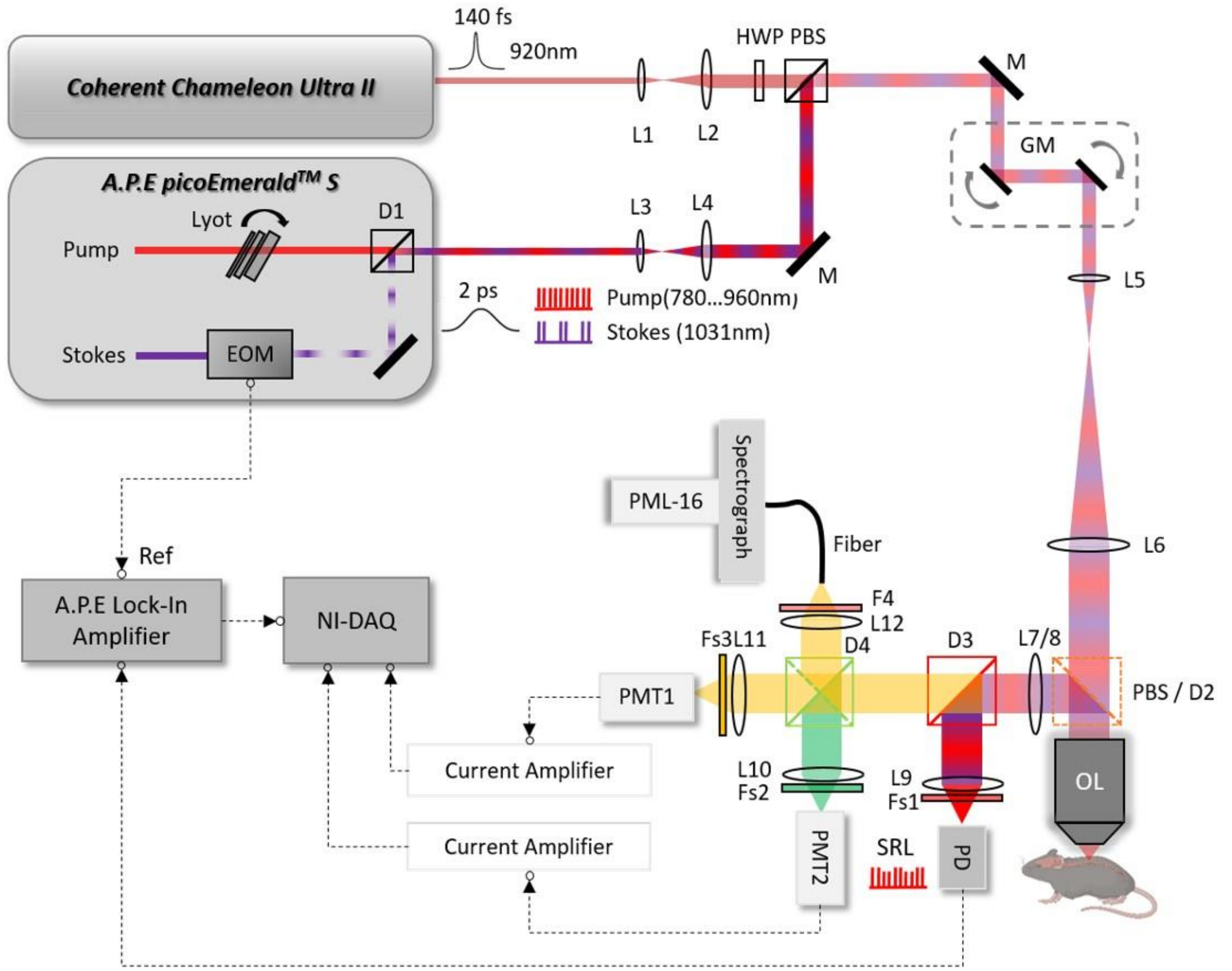


**Supplementary Figure 1. Schematic diagram of the multimodal NLO microscope system.** EOM: electro-optical modulator; HWP: half-wave plate; PBS: polarizing beam splitter; L1- L12: lenses; OL: objective lens; D1-D4: dichroic mirrors; Fs1-Fs3: filter sets; F4: filter; M: mirror; GM: galvo scanning mirrors; PD: photodiode; SRL: stimulated Raman loss; PMT1-2: photomultiplier tubes; PML-16: 16-channel photomultiplier tube; NI-DAQ: National Instruments data acquisition device.

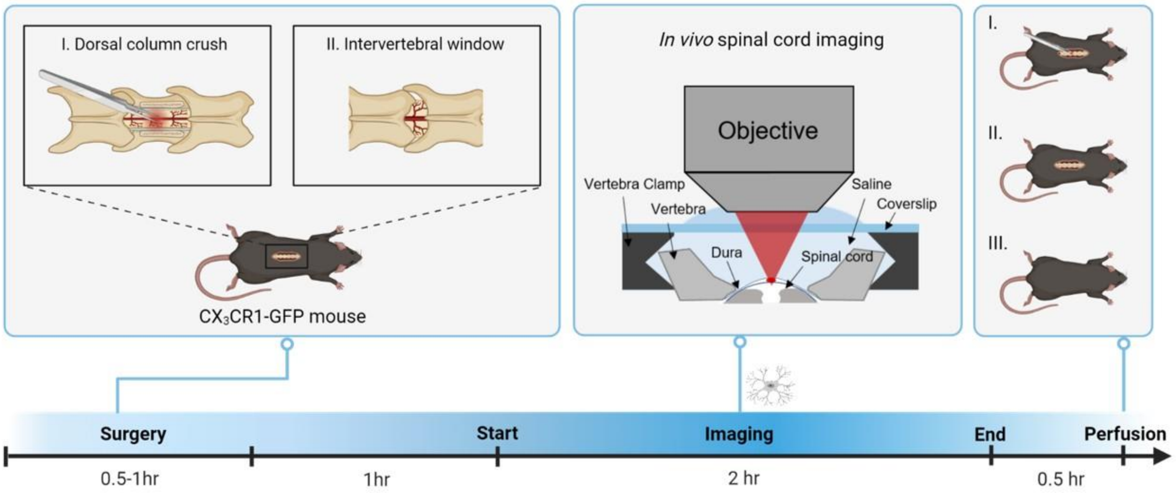


**Supplementary Figure 2**. **Experimental design for characterization of inflammation induced by the surgical preparations.** Group I, mice that underwent laminectomy and dorsal column crush (DCC); Group II, mice with spinal cord exposed in the intervertebral gap and only dura left above; Group III, mice that didn’t undergo surgery used as a negative control group (N.C.); 3 mice per group.

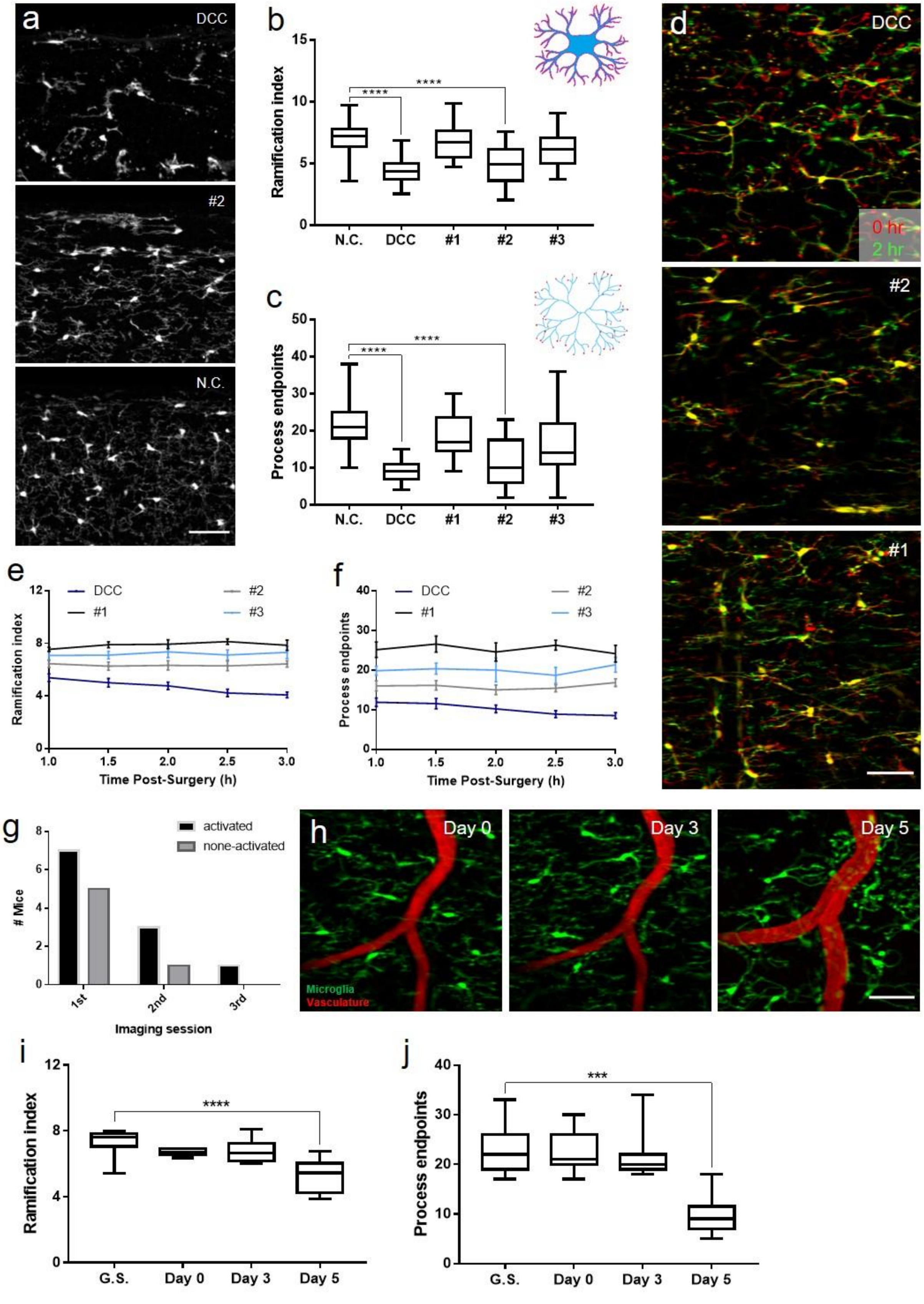
a
DCC
#2
N.C.
b
Ramification index
N.C.
DCC
#1
#2
#3
****
c
Process endpoints
d
0 hr
2 hr
e
f
Time Post-Surgery (h)
g
activated
none-activated
### Mice
1st
2nd
3rd
Imaging session
h
Day 0
Day 3
Day 5
Microglia
Vasculature
i
j
G.S.
***

12

**Supplementary Figure 3**. **Microglial behaviors under the conventional intervertebral window preparation** (a) Two-photon fluorescence images of 50-um-thick longitudinal spinal cord slices from the surgical and control site in mice expressing EGFP in microglia. #1-3, mouse#1-3 in the intervertebral window group; Scale bar, 50 $\mu$m. (b, c) Evaluation of microglia ramification index (b) and number of process endpoints (c) of spinal cord fixed slices from the three groups (Group N.C., DCC and Intervertebeal window); Kruskal-Wallis test: ****P<0.0001; n $\geq$ 20 measurements from 6-8 slices per mouse, three mice per group. The boxplots are shown with median, upper and lower quartiles and maximum and minimum values. (d) Representative *in vivo* superimposed images of microglia at an interval of two hours. Scale bar, 50 $\mu$m. (e, f) Changes of the microglia ramification index (e) and process endpoints (f) during two-hour *in vivo* imaging of intervertebral window and DCC group. n≥6 measurements per time point per mouse. Error bars, s.e.m. (g) Statistics of mice with microglia activation during longitudinal imaging through a conventional intervertebral window. (h) Longitudinal imaging of microglia (green) and vasculature (red) through a conventional intervertebral window at indicated times. Blood vessels were labeled with Texas Red dextran. Scale bar, 50 $\mu$m. (i, j) Ramification index (i) and process endpoints (j) of microglial cells shown in (h) and comparison with the gold standard (G.S.). Kruskal-Wallis test: ***P = 0.0002, ****P $\leq$ 0.0001; $n \geq 6$ microglial cells at each time were chosen for morphological quantification; The boxplots are shown with median, upper and lower quartiles and maximum and minimum values.

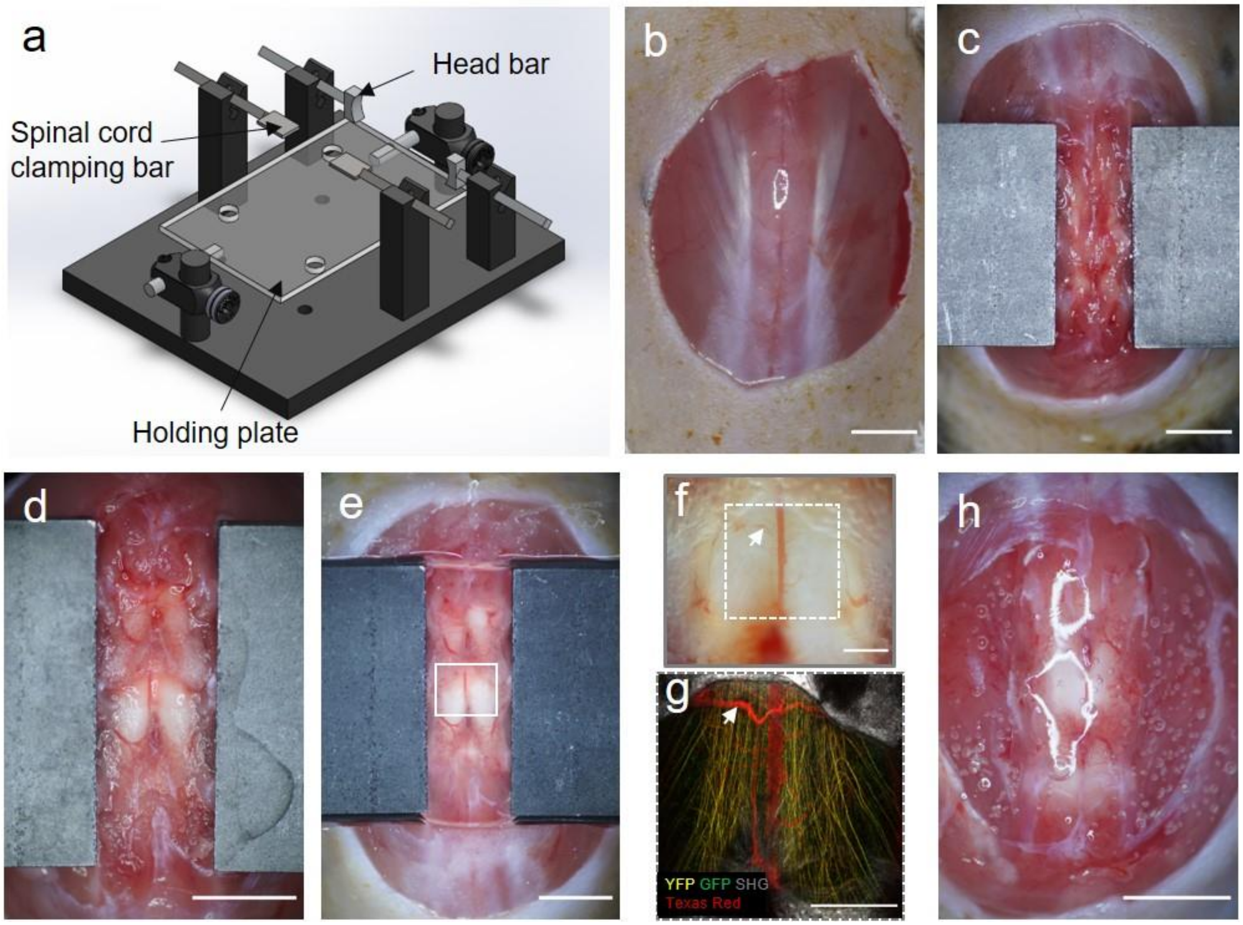


**Supplementary Figure 4. Surgical preparation of intervertebral LF window.** (a) A custom-designed stage for spinal cord stabilization during surgery and imaging. The stage consists of two spinal cord clamping bars to secure the spinal column. The two head bars attached with TACK-IT (FABER-CASTELL) are used to stabilize the mouse's head by holding its skull with slight pressure, which helps reduce motion artifacts caused by breathing. The holding plate with adjustable height is used to hold the mouse's body and allows more space for chest movement during breathing by moving it away from the mouse. (b-h) The procedure of the first surgical preparation of the LF window. (b) After hair removal and disinfecting the skin by Iodine solution, a small (~1.5cm) midline incision of the skin was made over the T11-T13 vertebrae and the skin was partly cut to expose the dorsal tissue. Scale bar, 3 mm. (c) Muscles as well as tendons both on both the top and sides were severed to clean the bone so that the spine can be held stably by clamping the vertebra from both sides. Scale bar, 3 mm. (d) Tissues attached to the vertebra were further resected.

Muscle tissues and tendons in the cleft between the vertebra arcs T12 and T13 were removed to expose the intervertebral window. Ligamentum flavum was left above the spinal cord dorsal surface. Additional care should be taken to avoid direct contact with the window surface. Scale bar, 3 mm. (e) A coverslip (22mm×22mm) was placed on the clamping bar and the interspace between the coverslip and cord was filled with saline/Iodixanol using a syringe. Scale bar, 3 mm. (f) Magnified bright-field image in the box region of (e). Blood vessels (white arrow) in the epidural space can be visualized in the bright-field image. Scale bar, 500$\mu$m. (g) Maximal projection of the TPEF and SHG image stack of the dashed box region in (f). The same blood vessel in the epidural space is also indicated by the white arrow. Blood vessels were labeled with Texas Red dextran by retro-orbital intravenous injection. Scale bar, 500 $\mu$m. (h) After imaging, liquid Kwik-Sil (World Precision Instruments) was applied on top of the surgical area covering the window for protection. It takes ~3min for Kwik-Sil to cure to a gel. The skin was then sutured and covered with burn cream (Betadine). Scale bar, 3 mm.

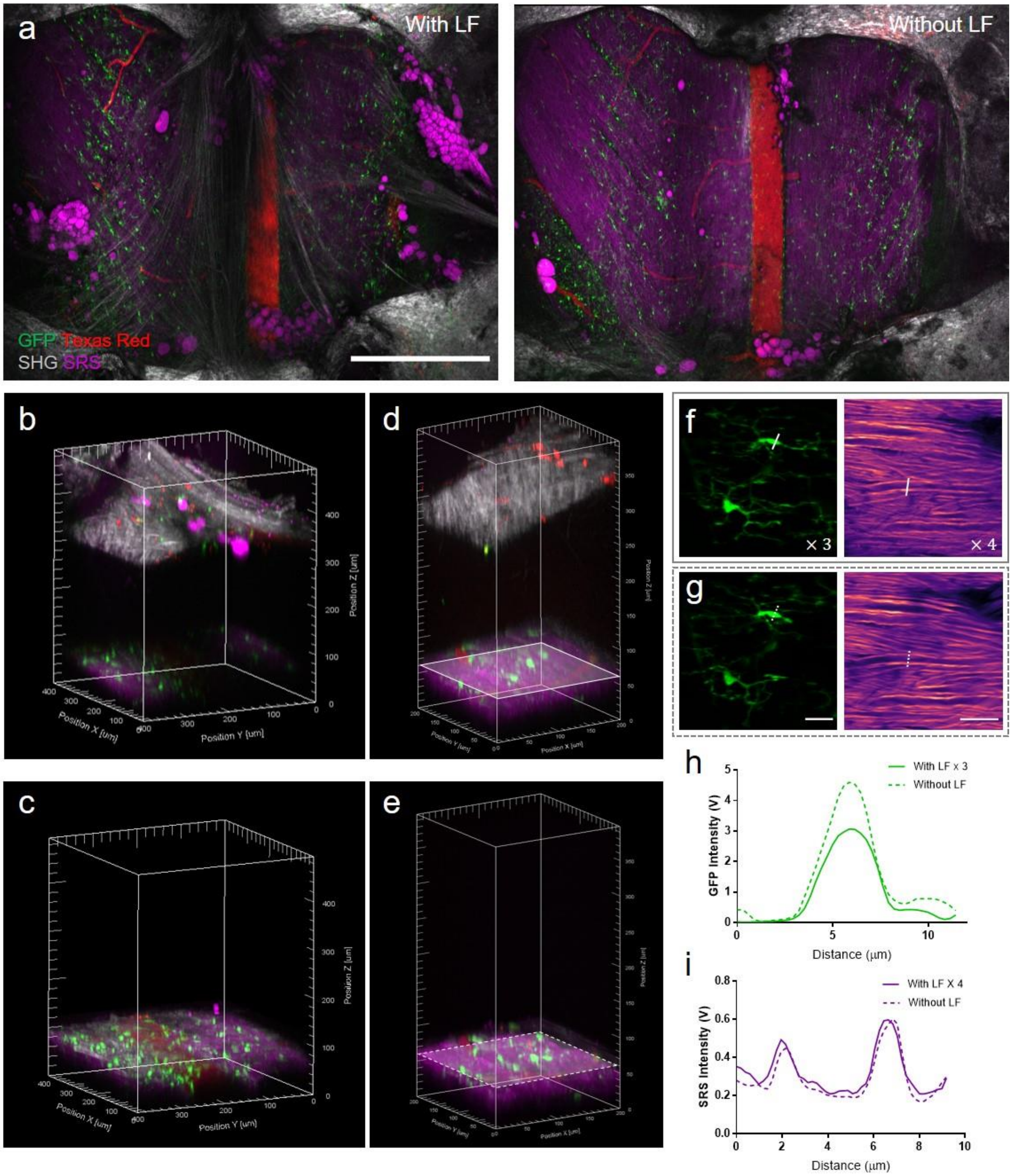


**Supplementary Figure 5**. **Comparison of LF window with the conventional one without LF for spinal cord imaging.** (a) Projection of an *in vivo* multimodal image stack of the intervertebral window before and after removal of ligamentum flavum. Scale bar, 500 $\mu$m. (b-e) 3D reconstruction of the multimodal imaging stacks at the location where spinal cord imaging is most affected by the existence of ligamentum flavum (b,c) and where imaging is less affected (d,e). Spinal cord *in vivo* imaging is compared at the same location before (b,d) and after (c,e) removal of ligamentum flavum. In (a-e): Green: GFP labeled microglia; Red, blood vessels and immune cells labeled with Texas Red dextran; Gray, second harmonic generation (SHG) signals of collagen and other connective tissues; Magenta, stimulated Raman scattering (SRS) signals of adipose tissue and myelin. (f, g) Maximal projection of TPEF microglia images (left) and SRS myelin images (right) at the same location of (d-e) before (f) and after (g) ligamentum flavum removal. Scale bar, 20 $\mu$m. Images in (f) and (g) are normalized to the same value and images in (f) were enhanced 3 and 4 fold digitally as indicated for better visualization. (h, i) Signal profile along the solid and dashed lines in (f) and (g) for a comparison of the fluorescence and SRS intensity before and after ligamentum flavum removal. SRS images are taken at Raman shift of 2863.5 $cm^{-1}$.

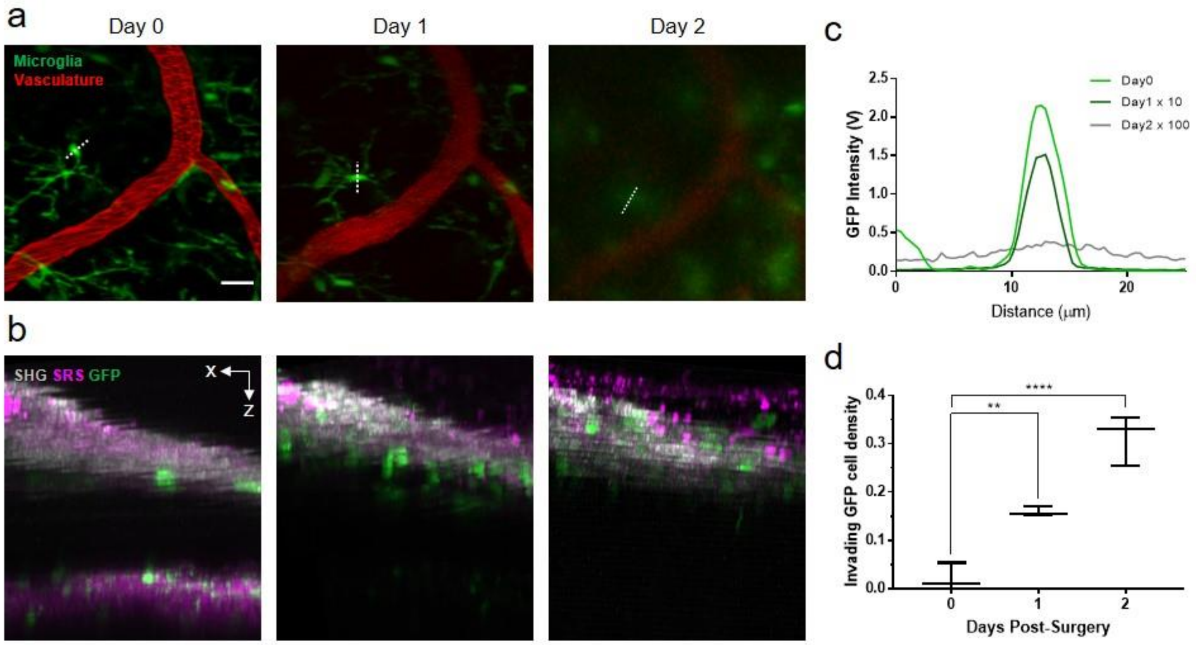


**Supplementary Figure 6**. **Longitudinal *in vivo* spinal cord imaging through LF window.** (a, b) Maximal x-y projection of microglia (green) and vasculature (red) images (a) under LF window and the corresponding x-z view (b) of the intervertebral window at indicated times. TPEF and SRS intensity in (a) and (b) are individually normalized for better visualization. Red, blood vessels labeled with Texas Red dextran; Gray, SHG signals of ligamentum flavum and meninges; Magenta, SRS signals of myelin and other tissues acquired at Raman shift of 2863.5 cm-1. Scale bar, 20 $\mu$m. (c) Signal profile along the dashed lines in (a) for a comparison of the fluorescence intensity at indicated times. (d) Density of the invading GFP cells in the same imaging volume of $300\mu m \times 300\ \mu m \times 100\ \mu m$ increases with time. One-way ANOVA: **P = 0.005, ****P≤0.0001. 1-2 ROIs measured for each mouse, three mice for each time point.

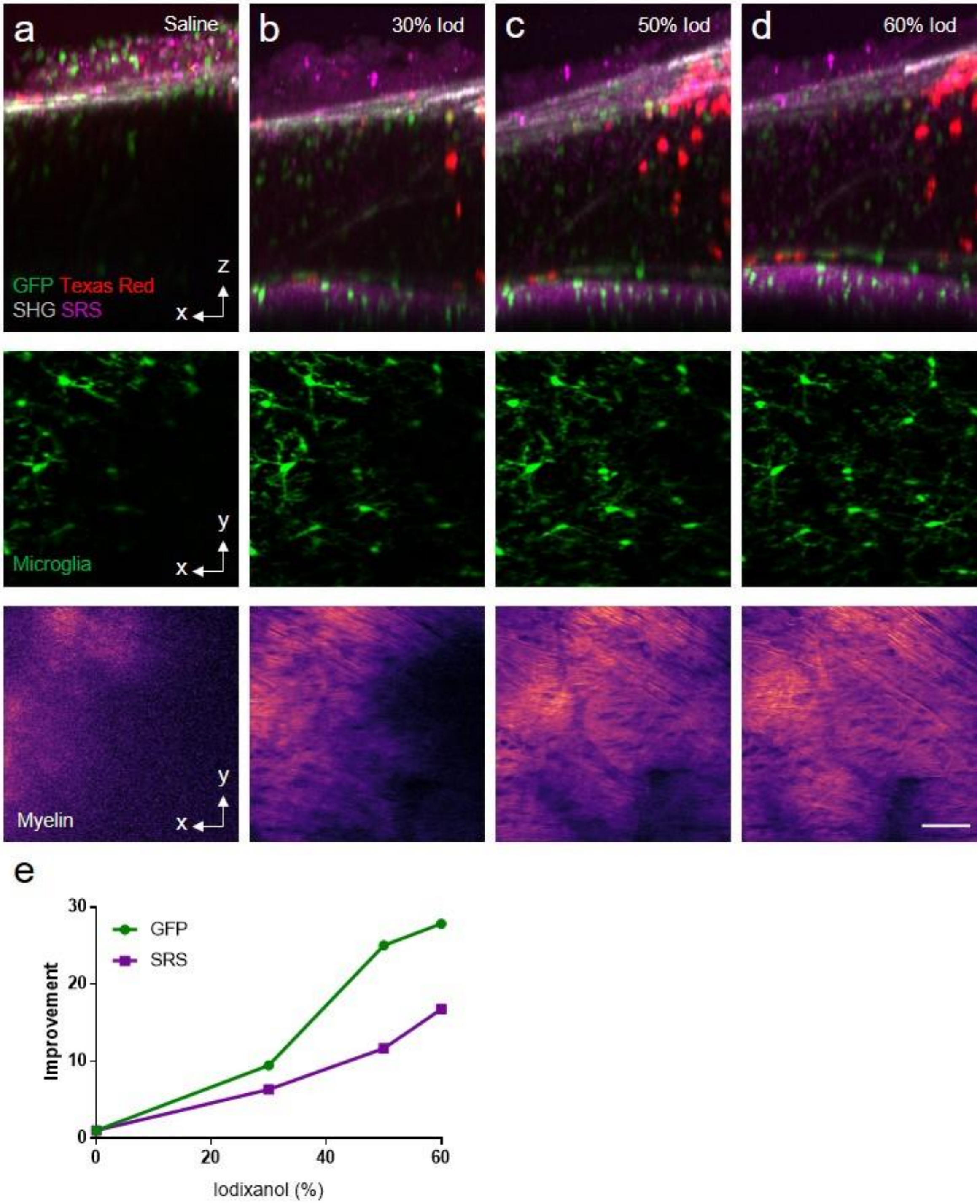


**Supplementary Figure 7. Effect of optical clearance as a function of Iodixanol concentration.** (a-d) Maximal x-z projection of a multimodal image of the intervertebral window and the maximal x-y projection image of microglia and myelin under the application of saline (a), 30% (b), 50% (c) and 60% (d) Iodixanol.

Green: GFP labeled microglia and other immune cells; Red, blood vessels and immune cells labeled with Texas Red dextran; Gray, SHG signals of ligamentum flavum and meninges; Magenta, SRS signals of myelin and other tissues acquired at Raman shift of 2863.5 $cm^{-1}$. Scale bar, 50 $\mu$m. (e) Quantification of the improvement in the microglia GFP and myelin SRS signal obtained by using Iodixanol at different concentrations.

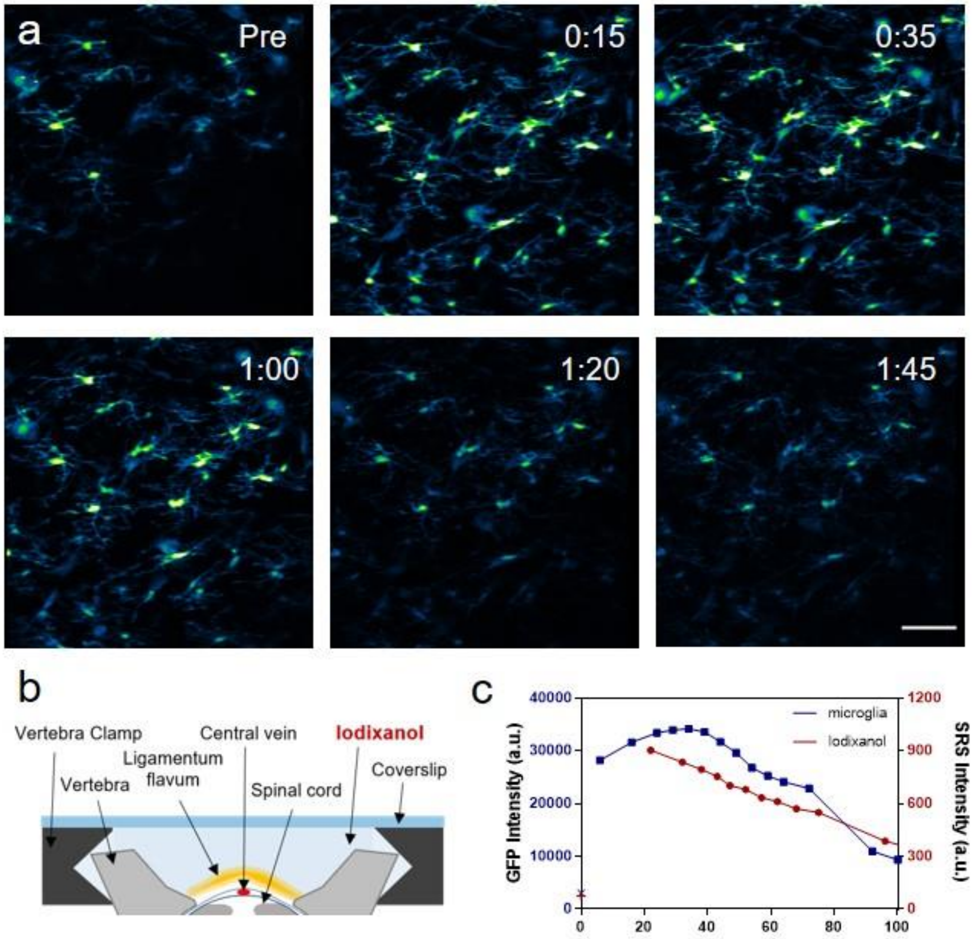


**Supplementary Figure 8. Iodixanol washout in LF window as a function of time.** (a) Maximal projection of microglia TPEF image stack at the indicated time before and after Iodixanol application. Images are normalized to the same value. Time is presented as hr:min. Scale bar, 50 $\mu$m. (b) Schematic diagram of the intervertebral window setup for optical imaging. Iodixanol fills the space between the spinal cord and coverslip. (c) GFP intensity of microglia and SRS intensity of Iodixanol as functions of time after the application of Iodixanol. Signals before Iodixanol application are denoted as cross symbols at time 0. GFP intensity is the average intensity of the brightest 0.1% pixels in the microglia maximal projection images. Iodixanol SRS images are taken 10 $\mu$m above the ligamentum flavum at Raman shift of 2943 $cm^{-1}$, corresponding to its Raman peak in the C-H region. SRS intensity is calculated as the average intensity of the image.

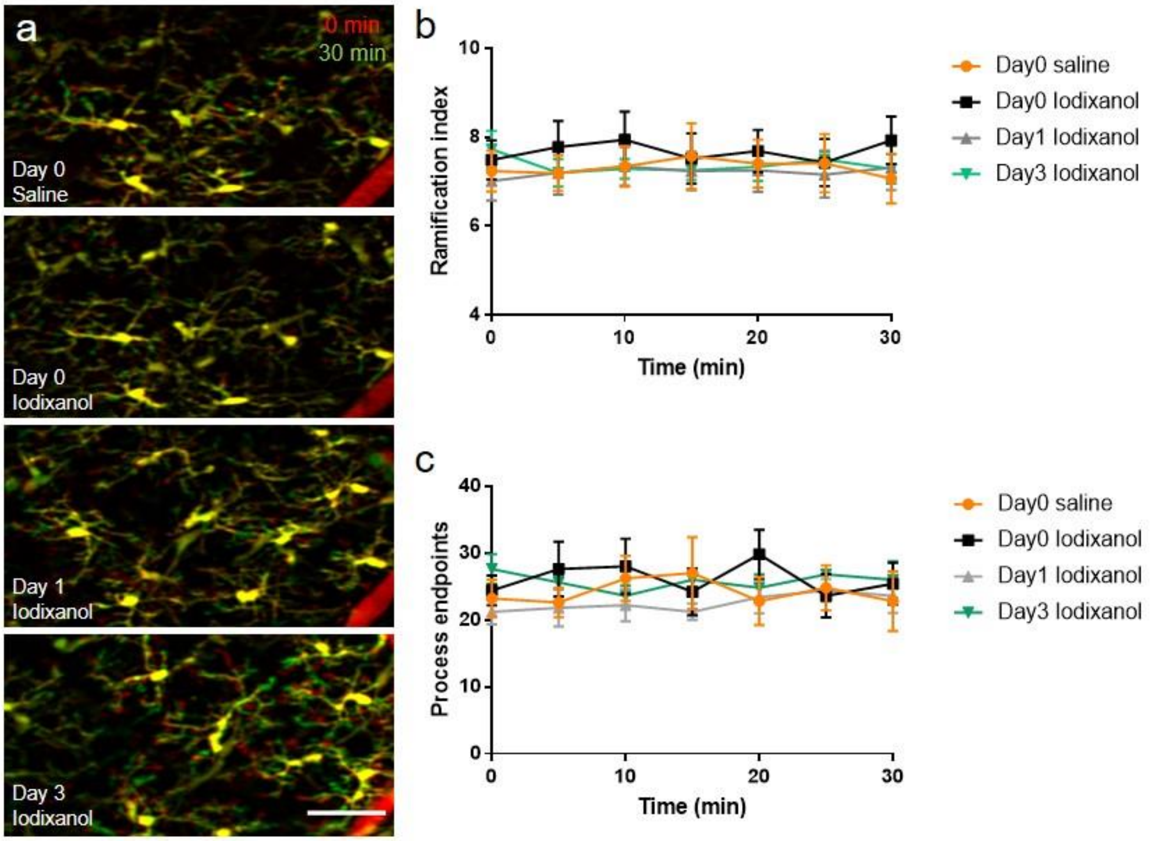


**Supplementary Figure 9. Effect of Iodixanol administration on microglial morphology as a function of time.** (a) Superimposed images of microglia at the same ROI at intervals of 30 min, showing process movement after saline/Iodixanol application at the indicated time. Blood vessels (red) labeled with Texas Red dextran are used as landmarks to navigate the same ROI. Scale bar, 50 $\mu$m. (b, c) Progress over time of the ramification index and process endpoints. Two mice for each time point. For each mouse, $n \geq 5$ microglial cells from the same ROI are selected for analysis. Error bar, s.e.m.

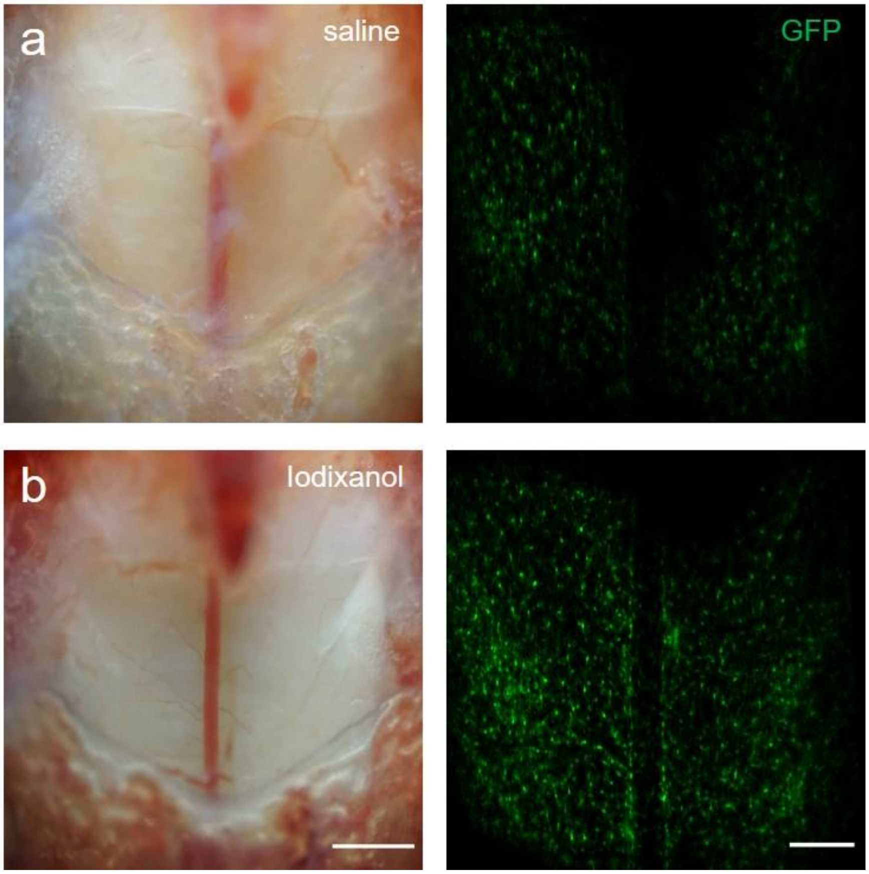


**Supplementary Figure 10. The improvement of intervertebral window clarity by optical clearing on day 0.** (a, b) Bright-field image and maximal projection of microglia TPEF image stack before (a) and after (b) optical clearing with 60% Iodixanol (w/v). Scale bar, 500 $\mu$m (bright-field), 200 $\mu$m (TPEF).

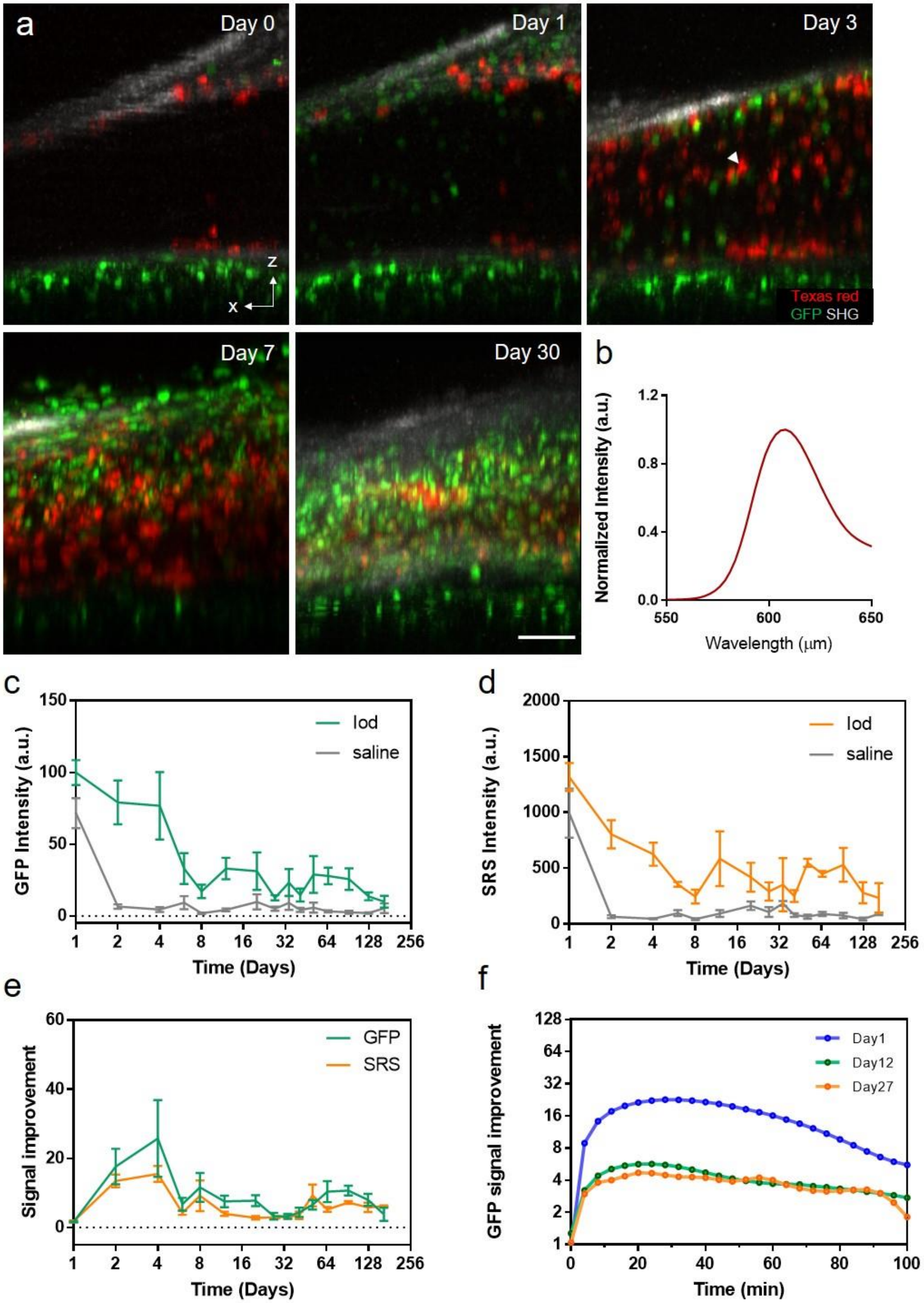
a
Day 0
Day 1
Day 3
z
x
Texas red
GFP SHG
Day 7
Day 30
b
Normalized Intensity (a.u.)
1.2
0.8
0.4
0.0
550
600
650
Wavelength (μm)
c
GFP Intensity (a.u.)
150
100
50
0
1
2
4
8
16
32
64
128
256
Time (Days)
Iod
saline
d
SRS Intensity (a.u.)
2000
1500
1000
500
0
1
2
4
8
16
32
64
128
256
Time (Days)
Iod
saline
e
Signal improvement
60
40
20
0
1
2
4
8
16
32
64
128
256
Time (Days)
GFP
SRS
f
GFP signal improvement
128
64
32
16
8
4
2
1
0
20
40
60
80
100
Time (min)
Day1
Day12
Day27

**Supplementary Figure 11. The clearing effect of Iodixanol as a function of time** (a) Maximal x-z projection images of the intervertebral window immersed in Iodixanol at the indicated time. Red, blood vessels and invading, likely inflammatory cells labeled with Texas Red dextran; Green: GFP labeled microglia and other immune cells; Gray, SHG signals of connective tissues; Scale bar, 50 $\mu$m. (b) Spectral properties of the red fluorescence excited by 920 nm femtosecond laser, indicating the fluorescence of Texas Red. This strong red fluorescence was found in the perivascular space and layers above the spinal cord indicated by the white arrowhead in (a). (c,d) GFP intensity of microglia (c) and SRS intensity of myelin (d) as functions of time after the first surgical procedure. At each time, GFP and SRS intensity were compared before and after Iodixanol application. (e) Improvement of GFP and SRS intensity by Iodixanol optical clearing as functions of time. In (c-e), GFP intensity is the average intensity of the brightest 0.1% pixels in the microglia maximal projection images. SRS intensity is the average intensity of the brightest 50% pixels in myelin maximal projection images. The improvement in signal is calculated as the ratio of signal intensity with and without optical clearing. At each time, the GFP and SRS intensity from the same ROI of each mouse were used for analysis. One to two ROIs were selected for analysis for each mouse, in total 4 mice were included for analysis. Error bar, s.e.m. (f) GFP signal improvement as functions of time after a single application of Iodixanol at the indicated time after the first surgery. GFP intensity is the average intensity of the brightest 0.1% pixels in the microglia maximal projection images.

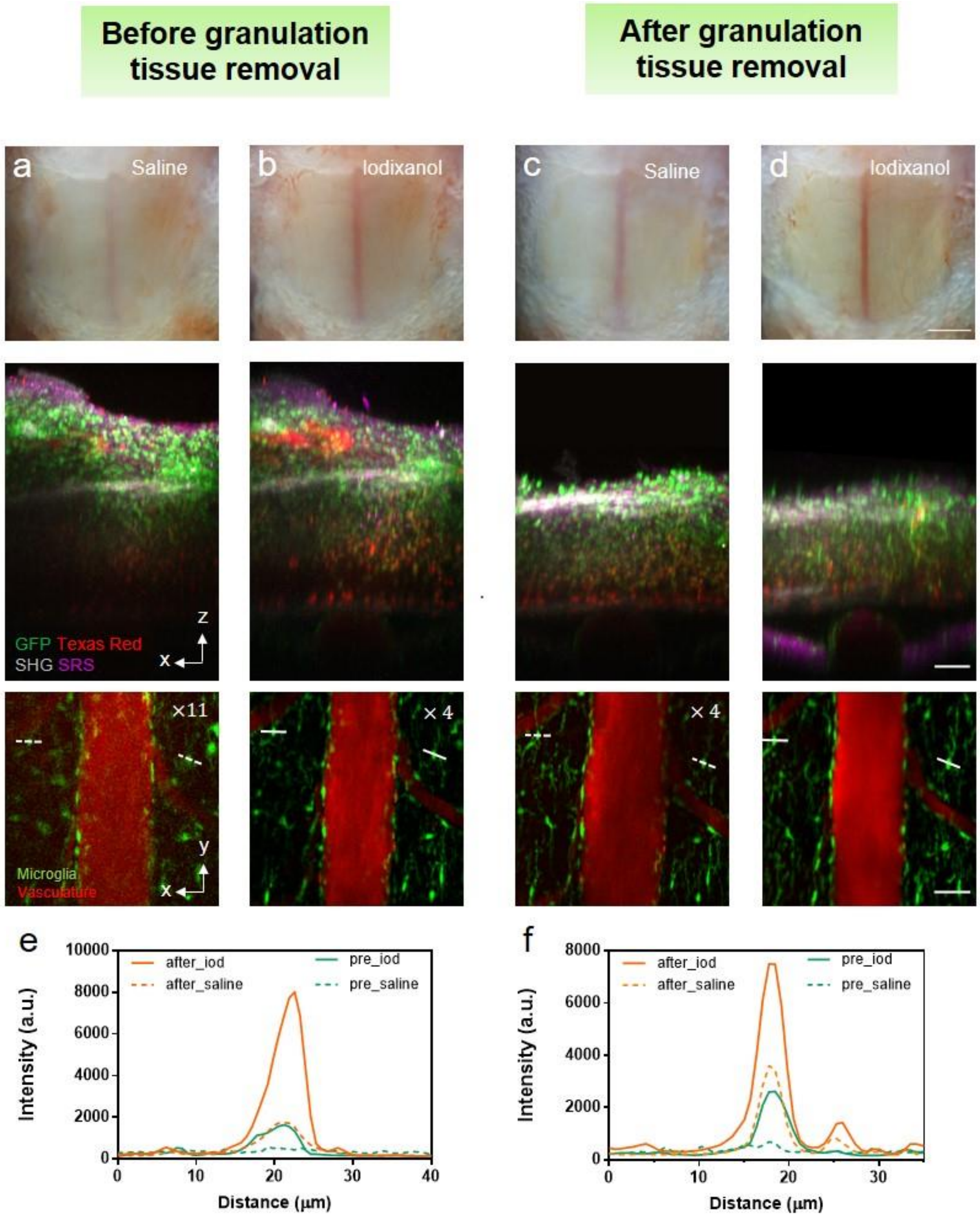


**Supplementary Figure 12. Improvement of window clarity by granulation tissue removal.** (a-d) Before and after removal of granulation tissue, comparison of the bright-field image, maximal x-z projection image of the intervertebral window and the corresponding x-y projection image of the spinal cord before and after Iodixanol application. x-y images of microglia and vasculature are normalized to the same value while images in (a-c) were enhanced digitally as indicated to improve visualization. Images were taken 19 days after the first surgery and 7 days after the last surgery. Red, blood vessels and immune-like cells labeled with Texas Red dextran; Green: GFP labeled microglia and other immune cells; Gray, SHG signals of connective tissues; Magenta, SRS signals of myelin and other tissues acquired at Raman shift of 2863.5 $cm^{-1}$. Scale bar, 500 $\mu$m for bright view image; 50 $\mu$m for x-z multimodal and x-y fluorescence images. (e-f) Intensity profile along the solid and dashed line in (a-d).

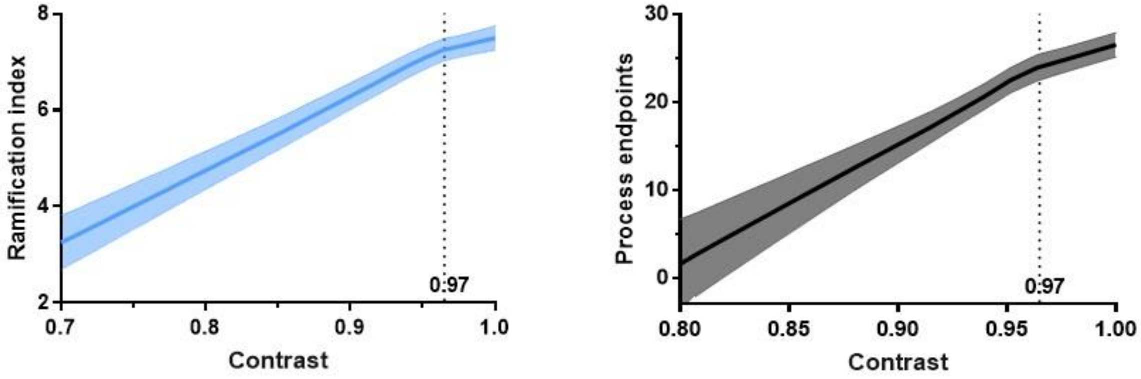


**Supplementary Figure 13. Ramification index and process endpoints of microglia as functions of fluorescence image contrast.** By applying Iodixanol with various concentrations to the intervertebral window, the relationship between the image contrast and morphological index was acquired for each microglia. All the curves of each microglia were averaged to get the functions of the image contrast and microglia morphological index. By curve fitting and slope analysis, image contrast of above 0.97 is used as the standard of microglial selection for morphological quantification. Error bar, s.e.m.

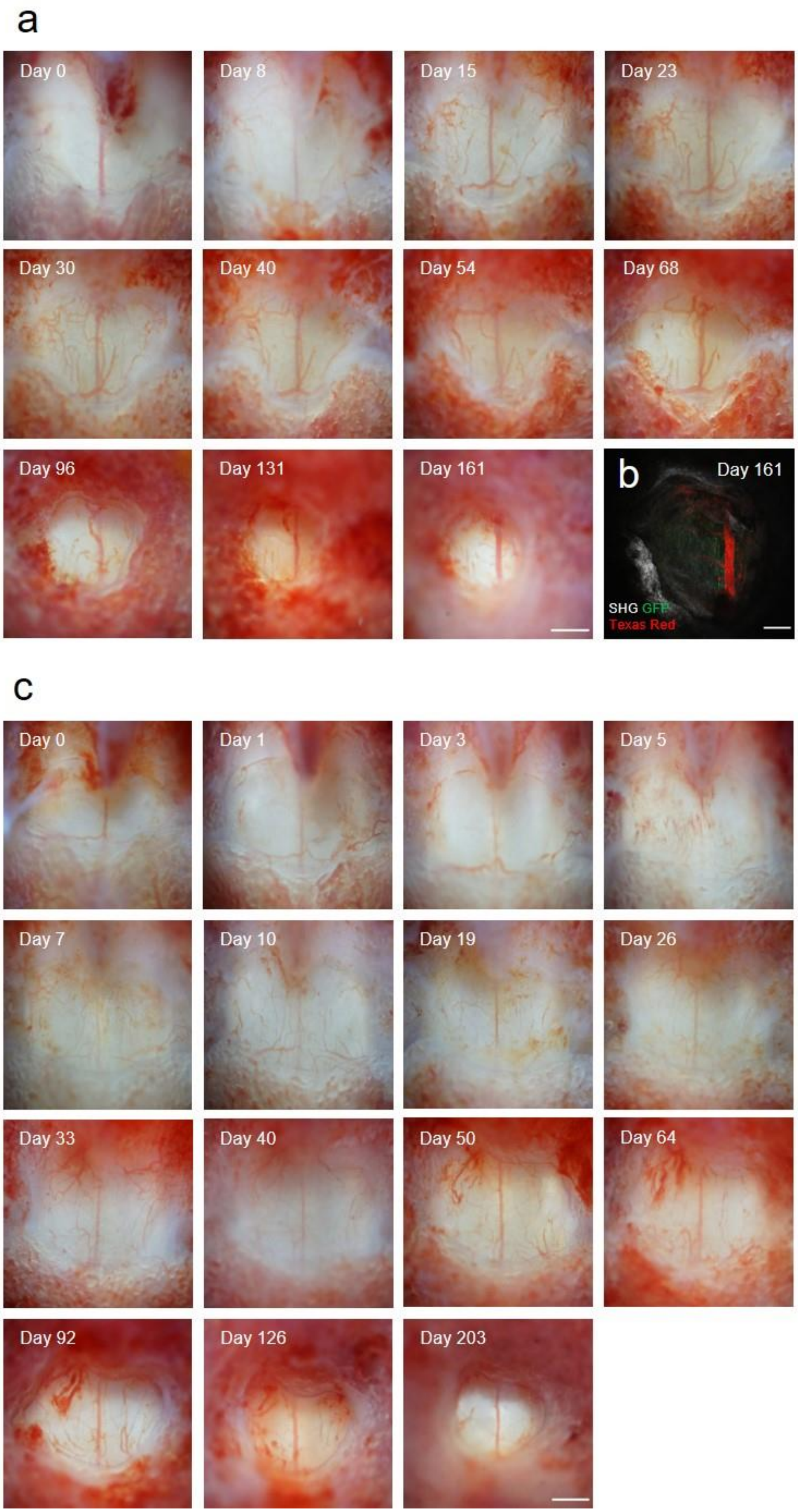
a
Day 0
Day 8
Day 15
Day 23
Day 30
Day 40
Day 54
Day 68
Day 96
Day 131
Day 161
b
Day 161
SHG GFP
Texas Red
c
Day 0
Day 1
Day 3
Day 5
Day 7
Day 10
Day 19
Day 26
Day 33
Day 40
Day 50
Day 64
Day 92
Day 126
Day 203

149

**Supplementary Figure 14. Decrease of FOV in LF window due to the growth of surrounding rigid tissue.** (a, c) Bright-field images of the intervertebral window from two mice at the indicated time after the first surgery, showing the decreased FOV after three months. Scale bar, 500 $\mu$m. (b) A two-photon image of the intervertebral window at day 161 shows strong SHG signal from the surrounding rigid tissue, suggesting the decrease of window FOV might be caused by the growth of the surrounding vertebra. Red, blood vessels labeled with Texas Red dextran; Green: GFP labeled microglia; Gray, SHG signals of connective tissues; Scale bar, 200 $\mu$m.

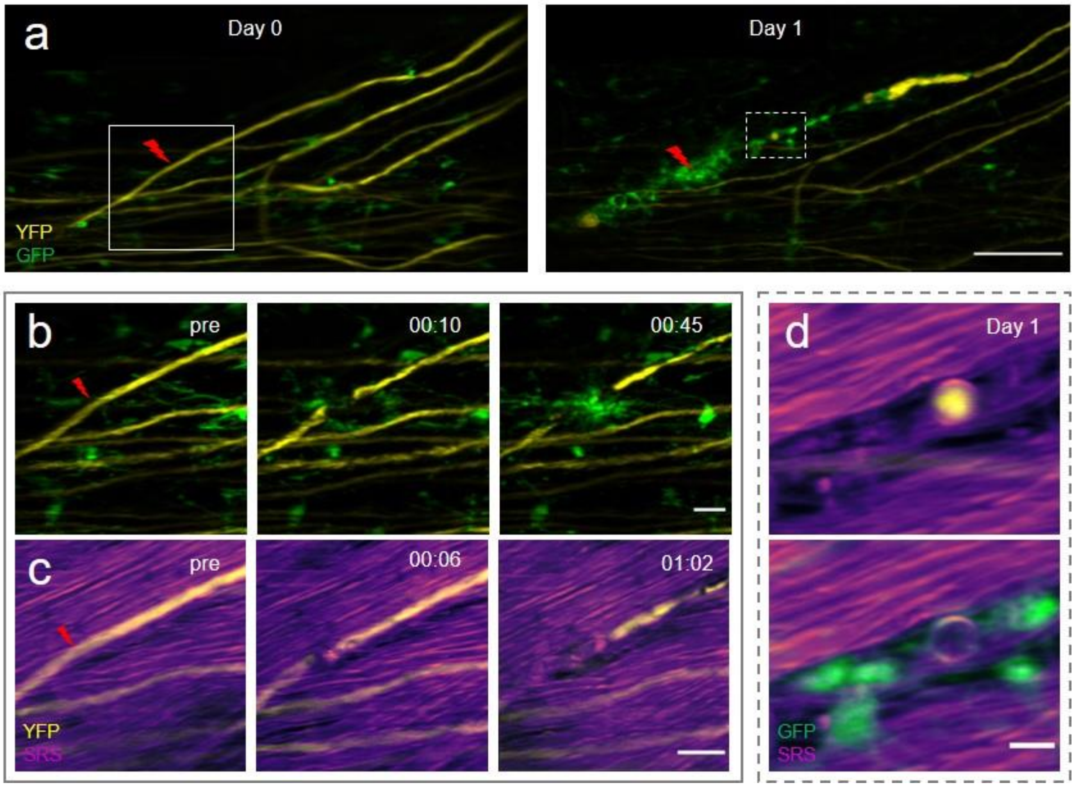


**Supplementary Figure 15. Multimodal NLO imaging of axonal degeneration on day 0 and day 1** (a) Maximal projections of TPEF image stacks of axons (yellow) and microglia (green) at the indicated times. The lightning bolt symbol indicates the lesion site. Scale bar, 100 $\mu$m. (b,c) Maximal projection images of microglia (green) and axons (yellow) with its surrounding myelin in the solid box region in (a) before and after laser axotomy. Time is presented as hr:min. Scale bar, 20 $\mu$m. (d) Single myelin SRS image merged with axon (yellow) and microglia (green) TPEF image on day1 in the dashed box region in (a). Scale bar, 10 $\mu$m. SRS images of myelin were taken at Raman shift of 2863.5 $cm^{-1}$.

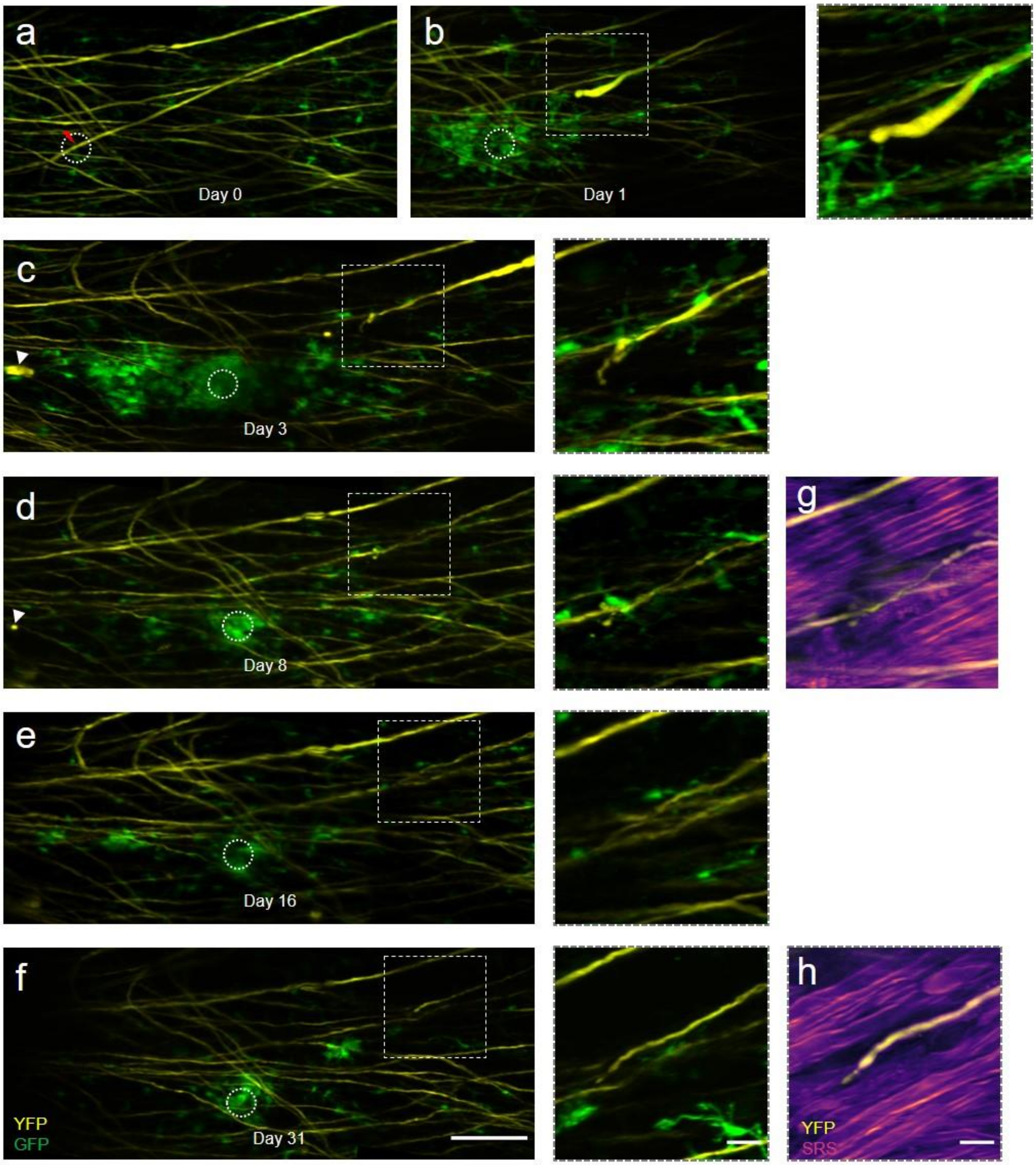


**Supplementary Figure 16. Multimodal NLO imaging of axonal degeneration after laser axotomy with a large lesion.** (a-f) Maximal projections of TPEF image stacks of axons (yellow) and microglia (green) at indicated times before and after laser axotomy. The lightning bolt symbol indicates the lesion site. The dashed circle indicates the size of injury. We determined the injury size based on observation of newly generated fluorescence or myelin disruption. Magnified images (b-f) of the box region are shown on the right. Arrowheads in (c) and (d) indicate the location of axonal debris. Scale bar, 100 $\mu$m, left; 20 $\mu$m, right. (g, h) Single SRS image of myelin merged with axon TPEF image (yellow) at day 8 and day 31. SRS images of myelin were taken at Raman shift of 2863.5 $cm^{-1}$. Scale bar, 20 $\mu$m.

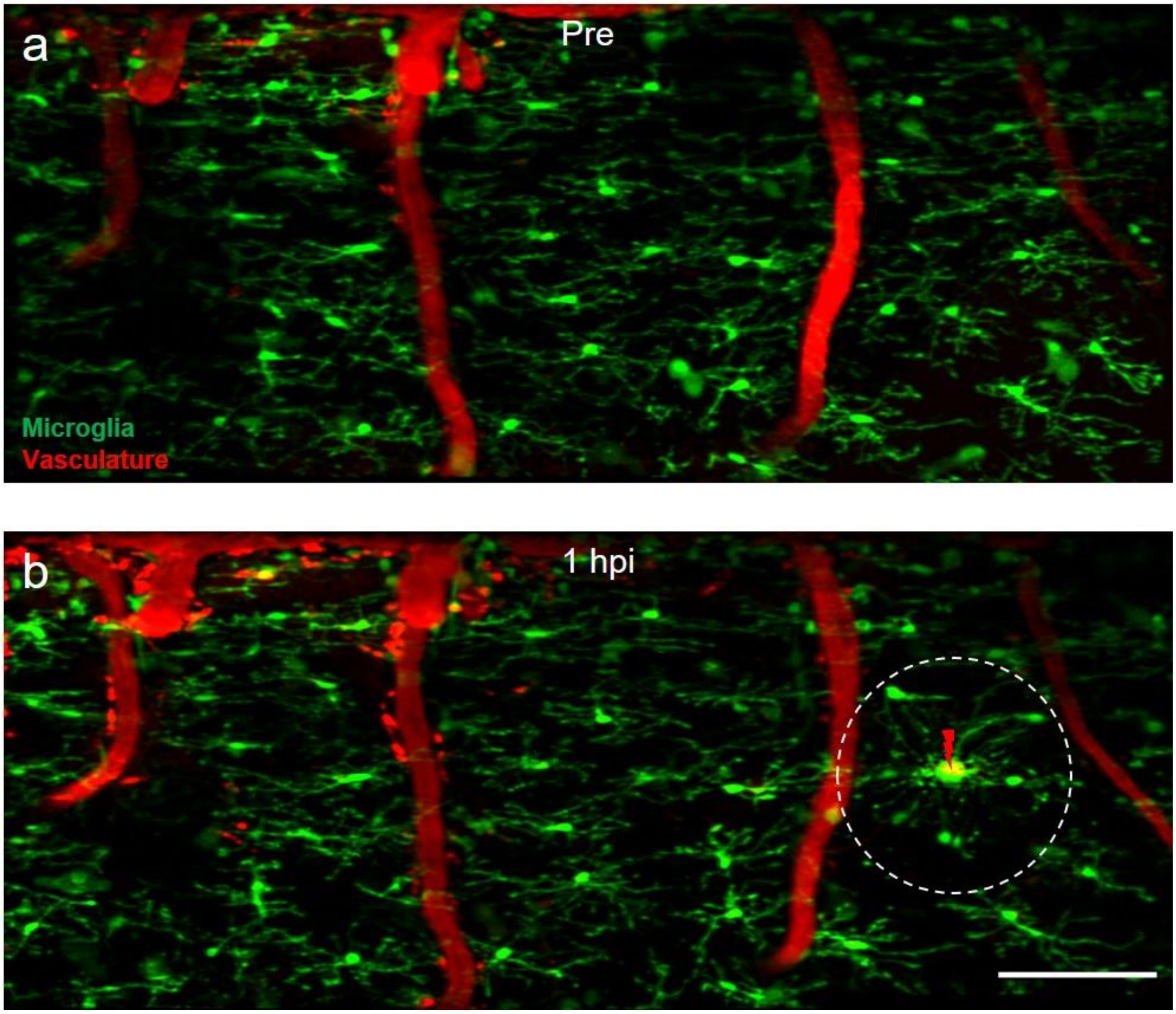


**Supplementary Figure 17. Precisely confined laser injury only affects microglial behavior in a limited region.** (a-b) Maximal projection image of microglia and vasculature before (a) and 1 hour after laser axotomy (b). The lightning bolt symbol indicates the lesion site. The dashed circle indicates the area in which the behavior of microglia is significantly affected by laser axotomy. The diameter of the dashed circle is 150 $\mu$m. Scale bar, 100 $\mu$m.

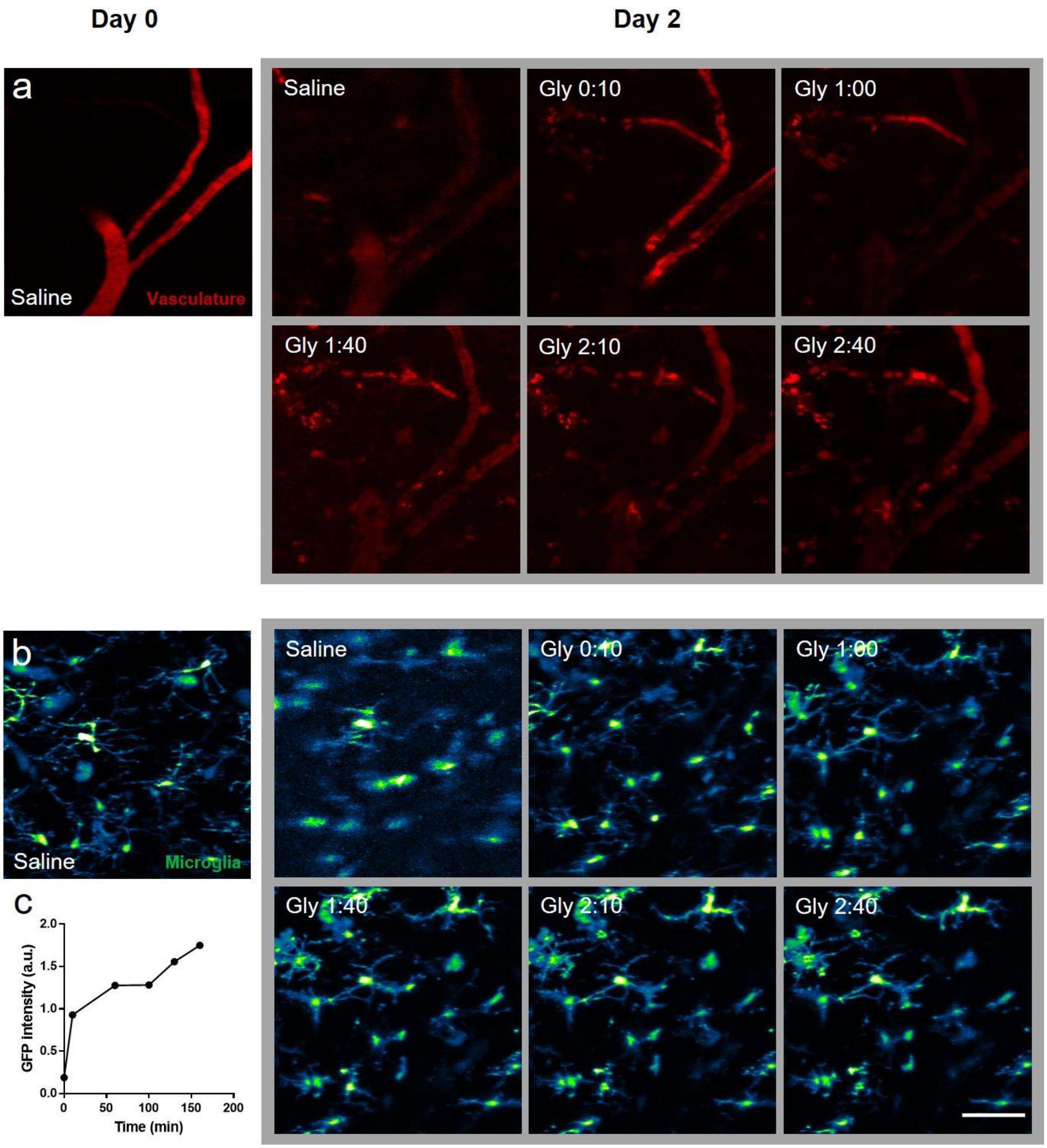


**Supplementary Figure 18. Optical clearing of LF window with glycerol** (a-b) Application of 70% v/v glycerol (G7757, Sigma-Aldrich) to the intervertebral window on day 2 helped to improve the image quality but induced vasculature disruption (a) and microglial activation (b), indicating glycerol toxicity to the spinal cord. Images of the same ROI taken on day 0 were used as a reference of normal microglial and vasculature morphology. The images presented are the maximal projection of the acquired TPEF image stacks. Blood vessels are labeled with Texas Red dextran. Time is presented as hr:min. scale bar, 50 $\mu$m. (c) Microglia GFP intensity as functions of time after glycerol application. GFP intensity is the average intensity of the brightest 0.1% pixels in the microglia maximal projection images. Application of glycerol can help to improve the image contrast and resolution under the intervertebral window but with toxic effects that have been reported in previous studies[1].

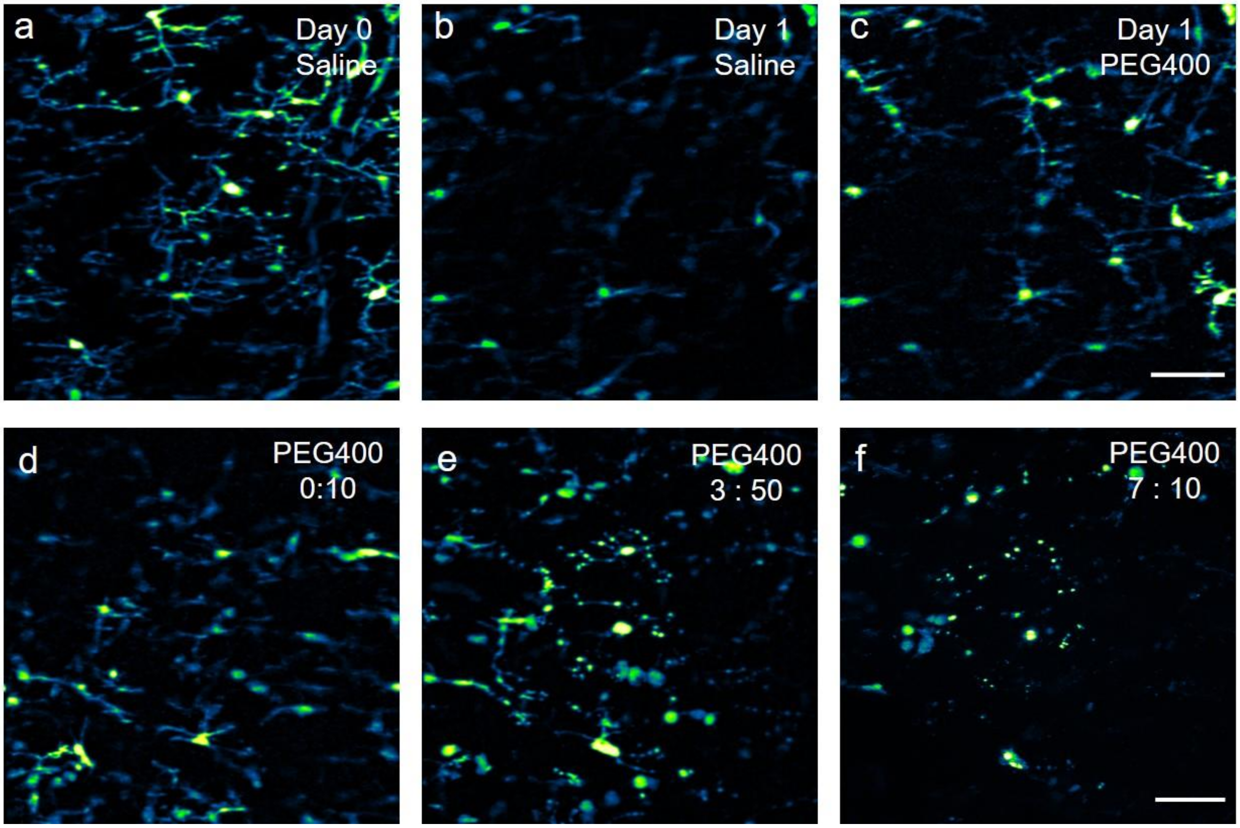


**Supplementary Figure 19. Optical clearing of LF window with PEG400** (a-c) Microglia maximal projection image on day 0 (a) and day 1 before (b) and 20 min after (c) application of 50% v/v PEG400 (P3265, Sigma-Aldrich). Image (b) and (c) are normalized to the same value. Scale bar, 50 $\mu$m. (d-e) Maximal projection image of microglia at the indicated time after PEG400 (50% v/v) application on day1. Enlarged soma size with beading processes of microglia was observed at the fourth hour after PEG400 application, which indicates the toxicity of PEG400 as an optical clearing agent for spinal cord imaging. Time is presented as hr:min. Scale bar, 50 $\mu$m.

#### REFERENCES

1. Zhu, D. *et al.* Short-term and long-term effects of optical clearing agents on blood vessels in chick chorioallantoic membrane. *JBO* **13**, 021106 (2008).
